# Drebrin mediates scar formation and astrocyte reactivity during brain injury by inducing RAB8 tubular endosomes

**DOI:** 10.1101/2020.06.04.134924

**Authors:** Juliane Schiweck, Kai Murk, Julia Ledderose, Agnieszka Münster-Wandowski, Imre Vida, Britta J. Eickholt

**Author notes:** these two authors contributed equally to this work (Juliane Schiweck, Kai Murk).

## Abstract

The brain of mammals lacks a significant ability to regenerate neurons and is thus particularly vulnerable. To protect the brain from injury and disease, damage control by astrocytes through astrogliosis and scar formation is vital. Here, we show that brain injury triggers an *ad hoc* upregulation of the actin-binding protein Drebrin (DBN) in astrocytes, which is essential for the formation and maintenance of glial scars *in vivo*. In turn, DBN loss leads to defective glial scar formation and excessive neurodegeneration following mild brain injuries. At the cellular level, DBN switches actin homeostasis from ARP2/3-dependent arrays to microtubule-compatible scaffolds and facilitates the formation of RAB8-positive membrane tubules. This injury-specific RAB8 membrane compartment serves as hub for the trafficking of surface proteins involved in astrogliosis and adhesive responses, such as β1-integrin. Our work identifies DBN as pathology-specific actin regulator, and establishes DBN-dependent membrane trafficking as crucial mechanism in protecting the brain from escalating damage following traumatic injuries.

## Main

The brain of higher mammals is vulnerable to traumatic injury and disease, but largely lacks the capacity to regenerate neurons and rarely regains function in damaged areas (Barker et al., 2018). This makes the containment of local pathological incidents critical to avoid the propagation of inflammation and neurodegeneration into the uninjured brain parenchyma.

Astrocytes are key players in protection of CNS tissue via a defense mechanism known as reactive astrogliosis. This process is associated with comprehensive changes in astrocyte morphology and function (Schiweck et al., 2018). Reactive astrocytes develop hypertrophies of soma and protrusions, while cells in proximity to large lesion sites polarize and extend particularly long processes. Dense arrays of such ‘palisades’ and hypertrophic astrocytes constitute glial scars, which enclose as physical barriers inflammatory cues and extravasating leucocytes, and limit the spread of damage (Faulkner et al., 2004; Frik et al., 2018). Glial scars are anatomically well described, but the molecular details controlling astrocyte reactivity are still poorly understood.

In the context of astrogliosis and scar formation, we studied drebrin (DBN), a cytoskeletal regulator, which stabilizes actin filaments by sidewise binding and by competing off other actin binding proteins (Grintsevich et al., 2010; Mikati et al., 2013). It is widely expressed, but has mainly been studied in neurons (Aoki et al., 2005). In cultured astrocytes, DBN has been described to maintain connexin 43 at the plasma membrane (Butkevich et al., 2004). Functional coupling of astrocytes into networks through gap junctions is essential to modulate neuronal transmission (He et al., 2020; Pannasch and Rouach, 2013). A deficit in DBN would therefore be expected to cause profound phenotypes *in vivo*. However, mouse models with acute or chronic DBN loss indicate non-essential functions of this actin binding protein during neuronal development, as well as during synaptic transmission and plasticity (Willmes et al., 2017). Instead, we identified DBN as important local safeguard mechanism in dendritic spines during conditions associated with increased oxidative stress (Kreis et al., 2019). Here, we characterize DBN as an injury-specific actin regulator in astrocytes, crucial for the wounding response and tissue protection in the brain. During injury, DBN provides a key switch to alter actin network homeostasis, which prepares the foundation for tubular endosomes, enabling polarized membrane trafficking of crucial surface receptors. In this role, DBN controls reactive astrogliosis required to form glial scars and to protect the susceptible CNS from traumatic brain injury.

## RESULTS

### DBN is an injury-induced protein in reactive astrocytes

To investigate endogenous DBN protein expression in astrocytes, we performed immunocytochemistry in 21 day *in vitro* (DIV) cortical cultures from mice containing both neurons and astrocytes. While we did not detect any DBN signals in astrocytes, high DBN levels were found in dendritic spines of neighboring neurons (Figure 1A). We exploited an *in vitro* scratch-wound model to induce astrocyte reactivity, which increased DBN expression with prominent labelling along the astrocyte processes (Figure 1A). In purified astrocyte cultures, where cells adopt polygonal morphologies, antibody labelling and western blot analyses identified low DBN expression (Figures 1B, S1). Following mechanical scratch injury, DBN protein levels were increased, reaching 3.9-fold of the baseline level 24 h post-injury (Figure 1B).

**Figure 1:**
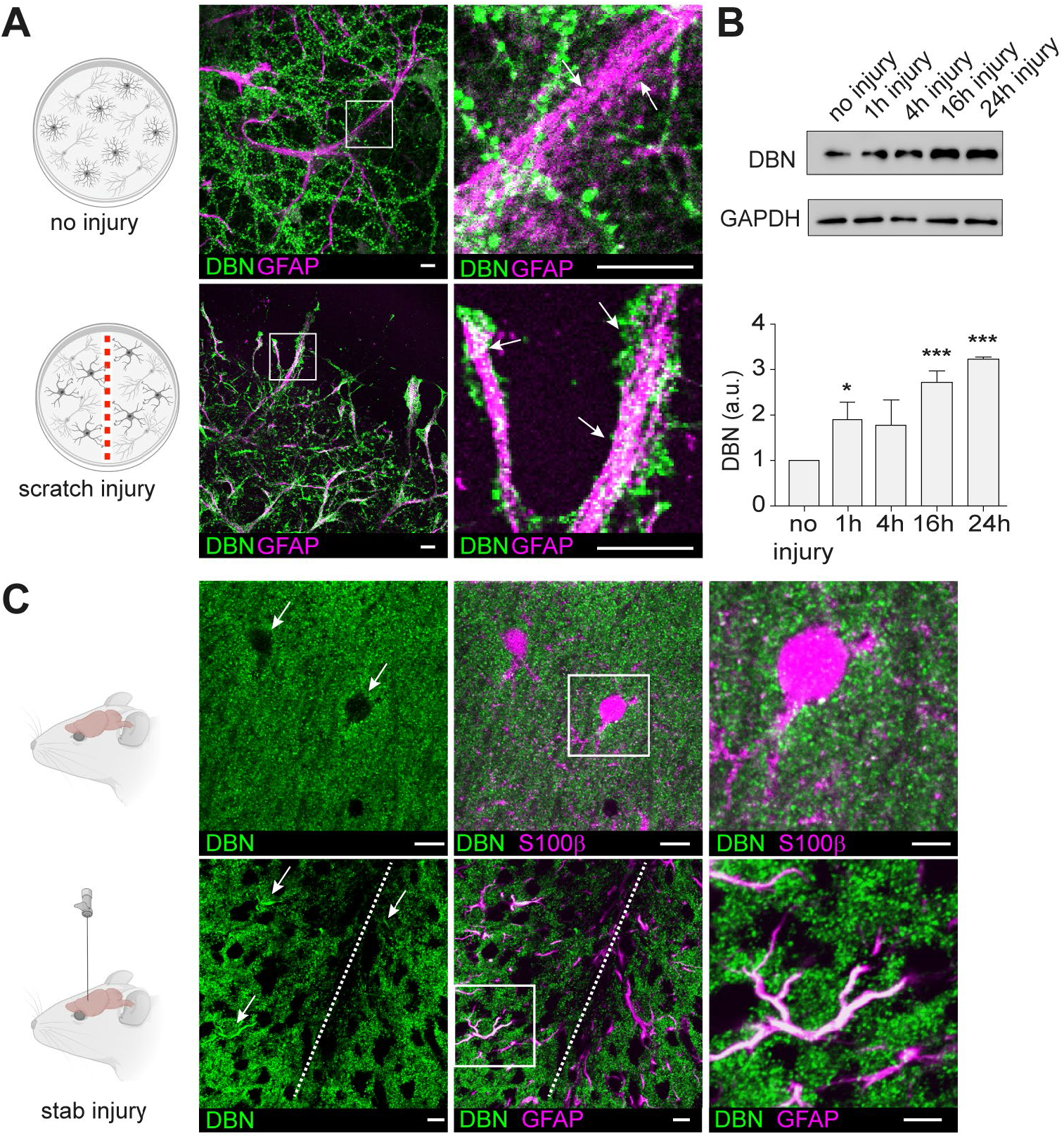
DBN protein levels are increased in response to injury. (A) DBN labeling (green) in mixed 21 DIV cortical cultures without (upper panels) or with mechanical scratch injury (lower panels). Square in left image is magnified on the right. Scale bars: 10 μm. (B) DBN expression analyses by western blotting in enriched astrocyte cultures without, or 1, 4, 16 and 24 h after scratch injury. Quantification of protein levels; n=3, \**P*<0.05, \*\*\**P*<0.001 (Student’s unpaired t-test). (C) Sections of P30 mouse brains without (upper panels) or with stab injury (lower panels) were labeled with anti-DBN (green) and anti-S100β (magenta). Scale bars: 10 μm. Magnifications on the right show no or high expression of DBN in S100ß+ or GFAP+ astrocytes in brains with or without stab injury, respectively. Scale bars: 20 μm.

In the mouse brain, DBN was distributed in distinct puncta in proximity to MAP2-positive dendrites, but not in S100ß+ astrocytes (Figure 1C). To induce reactive astrogliosis, we used an *in vivo* model for CNS trauma and inserted a needle unilaterally into the cortex. During this stab injury, numerous injury-induced GFAP+ astrocytes exhibited DBN protein expression throughout their polarizing processes (Figure 1C). In summary, our results demonstrate that CNS injury triggers an *ad hoc* upregulation of DBN in astrocytes *in vitro* and *in vivo*.

### Reactive astrocytes require DBN to form glia scars *in vivo*

To investigate if injury-induced upregulation of DBN protein is required during astrogliosis responses *in vivo*, we looked at its functional significance in DBN knockout mice during scar formation following cortical stab-injuries (Willmes et al., 2017). At 7 days post injury, the stab-wound triggered the formation of long palisading processes extending toward the injury site in 45% of GFAP-positive astrocytes in WT brains, in line with previous studies (Bardehle et al., 2013; Faulkner et al., 2004; Robel et al., 2011). In contrast, in *Dbn*^*−/−*^ mice, GFAP-positive astrocytes showed strong defects in polarization and in the formation of glial palisades barriers (Figure 2A).

**Figure 2:**
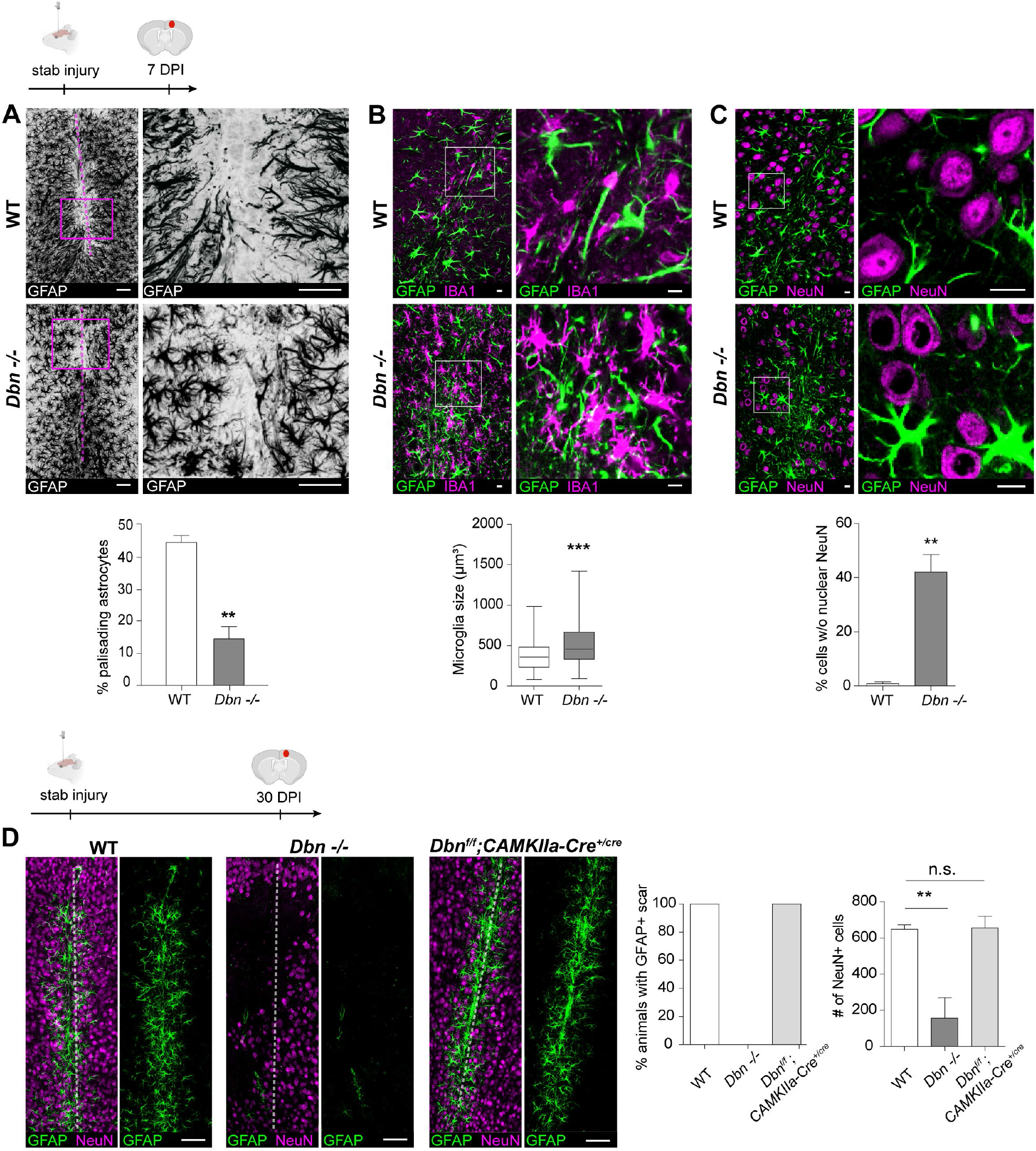
DBN controls glial scar formation and astrocyte reactivity *in vivo.* (A) GFAP+ astrocytes at core lesion sites in WT (upper panels) and *Dbn*^*−/−*^ (lower panels) brains, 7 days post stab-injury (DPI). Images show maximum projections of confocal planes at lesion sites. Magenta lines indicate the stab-injury from needle insertion. Magnifications show ‘palisading’ astrocytes extending towards injury sites. Scale bars: 100 μm. Quantification of palisading astrocytes in stab wounds 7 DPI. n=4 WT and 3 *Dbn*^*−/−*^ animals, **P<0.01 (Student’s unpaired t-test). (B) IBA1+ microglia (magenta) and GFAP+ reactive astrocytes (green) in the periphery of glial scars 7 DPI, in WT (upper panel) and *Dbn*^*−/−*^ brains (lower panel). Scale bars: 10 μm. Quantification of soma sizes of IBA1+ microglia. n=122-145 cells from 3 mice per condition, \*\*\**P*<0.001 (Student’s unpaired t-test). (C) NeuN+ neurons (magenta) and GFAP+ reactive astrocytes (green) in glial scars in WT (upper panel) and *Dbn*^*−/−*^ brains (lower panel) 7 DPI. Scale bars: 10 μm. Quantification shows neurons without nuclear NeuN in glial scars. n=5 WT and 3 *Dbn*^*−/−*^ mice, **P<0.01 (Student’s unpaired t-test). (D) NeuN+ neurons (magenta) and GFAP+ reactive astrocytes (green) in WT (left), *Dbn*^*−/−*^ (center) and *Dbn^fl/fl^:CAMK-Cre* brains (right), 30 DPI. Quantifications show percentage of animals with GFAP+ astrocytes (left graph) and NeuN+ cells (right graph) at lesion sites 30 days post-stab wounding. n= 4 WT and *Dbn*^*−/−*^ animals, 6 *Dbn^fl/fl^:CAMK-Cr*e animals, \*\**P*<0.01 (Student’s unpaired t-test).

Next we addressed whether the defect in scarring induced by DBN-loss affects the surrounding tissue. Microglia are highly sensitive sentinels in the brain, which become hypertrophic phagocytic cells after injury (Nimmerjahn et al., 2005). Whilst cultured microglia, as well as microglia in intact and injured brains do not express DBN (Figures S2A, B), they respond to injury with increased hypertrophy in *Dbn*^*−/−*^ brains when compared to WT brains (Figure 2B). This result indicates that stab-injury in *Dbn*^*−/−*^ brains exacerbates microglia responses, which likely reflects defective glial scarring.

We then asked if this defective glial scar development affects surrounding neurons. Loss of NeuN indicates degenerating neurons (Alekseeva et al., 2015), and its translocation from the nucleus to the cytosol identifies neurons exhibiting an initial stress response to pathologies (Lucas et al., 2014; Shandra et al., 2019; Wang et al., 2015; Wiley et al., 2016). Following stab injury, WT brains consistently maintained NeuN in the neuronal nuclei irrespective of their position relative to the injury site. In contrast, in *Dbn*^*−/−*^ brains, NeuN translocation from the nucleus to the cytosol was present in 42% of neurons (Figure 2C). As our disease model is considered as mild injury, we conclude that *Dbn*^*−/−*^ brains exhibit signs of increased brain damage 7 days post-injury.

### DBN provides long-term tissue protection after traumatic brain injury

To study the long-term outcome of DBN deficiency, we extended the stab-injury analyses to a later time point. At 30 days post-injury, WT mice maintained well-defined GFAP+ scars exhibiting typical signs of long-term thinning (Figure 2D) (Bardehle et al., 2013), whilst we no longer detected GFAP+ astrocytes in lesion sites of *Dbn*^*−/−*^ brains (Figure 2D). At the same time, we identified a substantial loss of NeuN+ cell bodies surrounding lesions in *Dbn*^*−/−*^, but not in WT brains (Figure 2D). These findings show long-term neurodegeneration upon cortical stab-injuries in DBN-deficient mice.

Finally, to exclude the possibility that neurodegeneration in *Dbn*^*−/−*^ brains following stab-injury is due to generally increased neuronal vulnerability rather than the observed defective astrocyte scarring, we analyzed the outcome of stab-injury in the brain of *Dbn^fl/fl^:CAMK-Cre^+/cre^* (*Dbn^fl/fl^:CAMK-Cre*) mice. These mice lose DBN expression in neurons, but not in astrocytes. Following cortical stab-injury, *Dbn^fl/fl^:CAMK-Cre* animals consistently show typical GFAP+ astrocytes in lesions, with no signs of NeuN-loss 30 days after stab-injury (Figure 2D). In summary, DBN is essential for astrocyte reactivity and damage containment during traumatic brain injury. DBN-loss perturbs injury-induced scar formation and suppresses maintenance of astrocyte reactivity, providing a causal link of DBN function during injury-induced reactive astrogliosis.

### DBN controls an injury-induced RAB8 membrane compartment

Towards identifying the DBN-dependent cellular mechanism in the injury setting, we employed the *in vitro* scratch injury of cultured astrocytes. Following injury, WT astrocytes polarized, and extended long processes into the injury site (Figure S3A). In contrast, the majority of *Dbn*^*−/−*^ astrocytes extended over smaller distances and frequently changed their orientation (Figure S3A; Movie 1), resulting in a significant decrease in wound closure (Figure S3B). During the wounding response, DBN, as well as overexpressed DBN-YFP only partially localized to prominent actin fibers and also decorated internal compartments consisting of vesicular and tubular structures (Figures S4A, B).

Given this subcellular distribution in astrocytes, together with a previous observation that identified DBN in association with internal membrane structures (Xu and Stamnes, 2006), we asked if DBN mediates injury responses by regulating membrane trafficking. Analyses of several trafficking compartments, including RAB5+, RAB7+ or RAB11+ endosomes showed no major differences in Dbn-/- astrocytes when compared to WT cells (Figure S5A). However, DBN-loss severely disturbed the distribution of RAB8 in astrocytes during mechanical injury. In WT astrocytes, using either GFP-RAB8A or an antibody that detects both RAB8A and RAB8B, we obtained signals of prominent RAB8+ tubular compartments adjacent to injury sites (Figure 3A, Figure S5A). No signals were detected in cells depleted for both RAB8A and RAB8B demonstrating specificity to the antibody (Figures S5B, C). In *Dbn*^*−/−*^ astrocytes, instead of marking internal tubules, RAB8 was mostly dispersed throughout the cytosol (Figure 3A).

**Figure 3:**
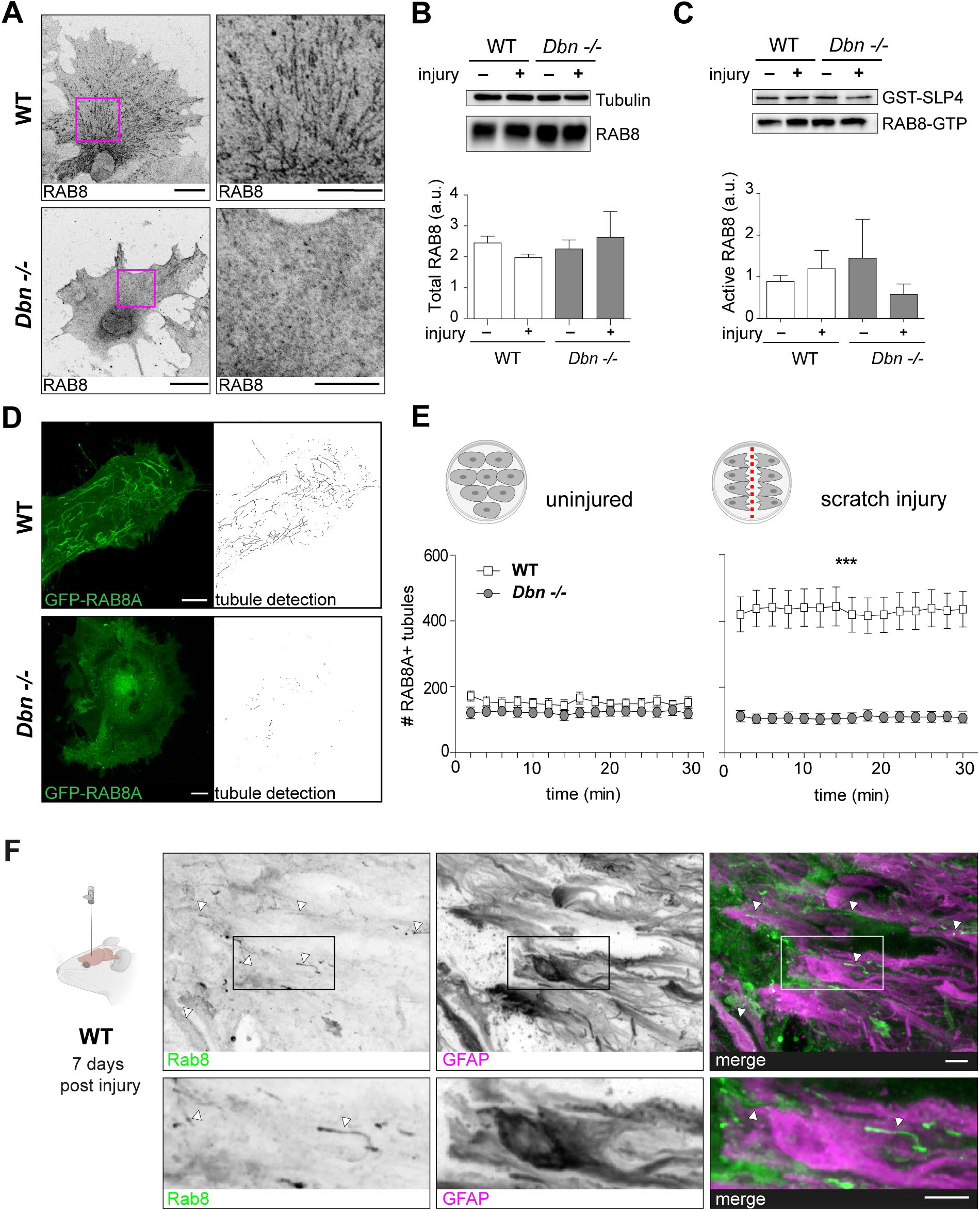
DBN controls the formation RAB8 tubular endosomes during injury in cultured astrocytes. (A) RAB8 labeling in WT (upper panels) and *Dbn*^*−/−*^ astrocytes (lower panels) 24 h after scratch injury *in vitro*. Scale bar: 10 μm. (B) Western blot analysis of RAB8 protein in WT and *Dbn*^*−/−*^ astrocytes without or 24 h after injury. Tubulin serves as loading control. Graph shows quantification of RAB8 protein levels in WT and *Dbn*^*−/−*^ astrocytes (n=6, Student’s unpaired t-test). (C) Western blot analysis of active RAB8-GTP isolated by GST-SLP4 pulldown from lysates obtained from WT and *Dbn*^*−/−*^ astrocytes without or 24 h after scratch injury (n=6, Student’s unpaired t-test). GST levels confirm equal amounts of SLP4 in pulldowns. Quantification shows GTP-bound RAB8 levels in WT and *Dbn*^*−/−*^ astrocytes (n=6, Student’s unpaired t-test). (D) GFP-RAB8A distribution in transfected WT (upper panels) and *Dbn*^*−/−*^ astrocytes (lower panels). GFP-RAB8A labelled compartments were detected automatically and are shown as skeletonized structures (black). Scale bars: 10 μm. (E) Quantification of GFP-RAB8A+ tubules in WT and *Dbn*^*−/−*^ astrocytes, uninjured (n=9-10) and 24 h after scratch injury (n=28-32, *P*<0.001; repeated measurements ANOVA). Cells were imaged for 30 min. (F) Presence of Rab8+ tubules (green) in GFAP+ astrocytes (magenta) at the lesion core following stab-injury in the brain. Tissue was labelled at 7 DPI. Arrowheads show Rab8 tubules. Scale bars: 10 μM.

Western blotting confirmed that RAB8 protein levels and GTPase activity in *Dbn*^*−/−*^ astrocytes were not altered when compared to WT, irrespective of injury (Figures 3B, C), indicating that DBN is required during the generation of the tubule compartment rather than influencing the activity of the GTPase. Analogous to endogenous RAB8, GFP-RAB8A highlighted prominent membrane tubules in WT astrocytes, while GFP-RAB8A tubules in *Dbn*^*−/−*^ were fragmented (Figure 3D, Movie 2). Quantification of tubules in uninjured astrocyte cultures revealed very few RAB8A-positive tubules in both WT and *Dbn*^*−/−*^ astrocytes. However, mechanical scratch injury increased the tubule number by 2.5-fold in WT astrocytes but not in *Dbn*^*−/−*^ astrocytes (Figure 3E), which indicates that tubular RAB8A endosomes form in response to injury and depend on the presence of DBN. We also detected Rab8+ tubules in polarizing astrocytes at lesion sites *in vivo* suggesting that these structures are also regulators of astrogliosis in the setting of brain injury and scarring responses *in vivo* (Figure 3F).

### DBN generates tubular endosomes by counteracting Arp2/3-dependent actin nucleation

Dynamic rearrangement of the actin cytoskeleton is a critical component in generating tubular membrane systems (Gautreau et al., 2014; Simonetti and Cullen, 2019). It relies on the capacity of actin filaments to polymerize and depolymerize, a characteristic that can be influenced by the ability of DBN to bind the lateral filament surface, to bundle actin filaments and to inhibit actin nucleation (Ginosyan et al., 2019; Grintsevich et al., 2010; Li et al., 2019). We analyzed whether defects of *Dbn*^*−/−*^ astrocytes in RAB8 compartments are linked to DBN and its function as actin regulator. In the first set of experiments, we co-transfected astrocyte cultures isolated from *Dbn*^*−/−*^ mice with DBN-YFP (or control YFP) and labelled RAB8A using red fluorescent mRuby-RAB8A. Following *in vitro* injury, DBN-YFP expression, but not YFP, rescued the ability of *Dbn*^*−/−*^ astrocytes to form membrane tubules (Figure 4A). Next, we employed pharmacological approaches to address whether defective actin dynamics disrupts Rab8a tubules in *Dbn*^*−/−*^ astrocytes. Low concentrations of Cytochalasin D (100 nM) interfere with the nucleation of new actin filaments whilst leaving existing filaments unaffected (Kim et al., 2010). During scratch-injury in *Dbn*^*−/−*^ reactive astrocytes, this Cytochalasin D concentration re-established RAB8A-positive membrane tubules (Figure 4B, Movie 3), suggesting that DBN induces tubule formation by antagonizing the cellular actin nucleation machinery. In this model, loss of DBN would increase actin nucleation. To test this idea, we applied the ARP2/3 inhibitor CK-666 or the formin-inhibiting molecule SMIFH2 (Nolen et al., 2009; Rizvi et al., 2009) to GFP-RAB8A expressing *Dbn*^*−/−*^ astrocytes. Whilst inhibition of the ARP2/3 nucleation machinery by CK-666 reinstalled the ability of *Dbn*^*−/−*^ astrocytes to form RAB8A tubules, the formin inhibitor did not (Figures 4B, Movie 3). Thus, DBN stabilizes actin filaments by mechanisms involving suppression of ARP2/3-dependent actin nucleation, a process essential for the injury-induced formation of the tubular RAB8 compartment.

**Figure 4:**
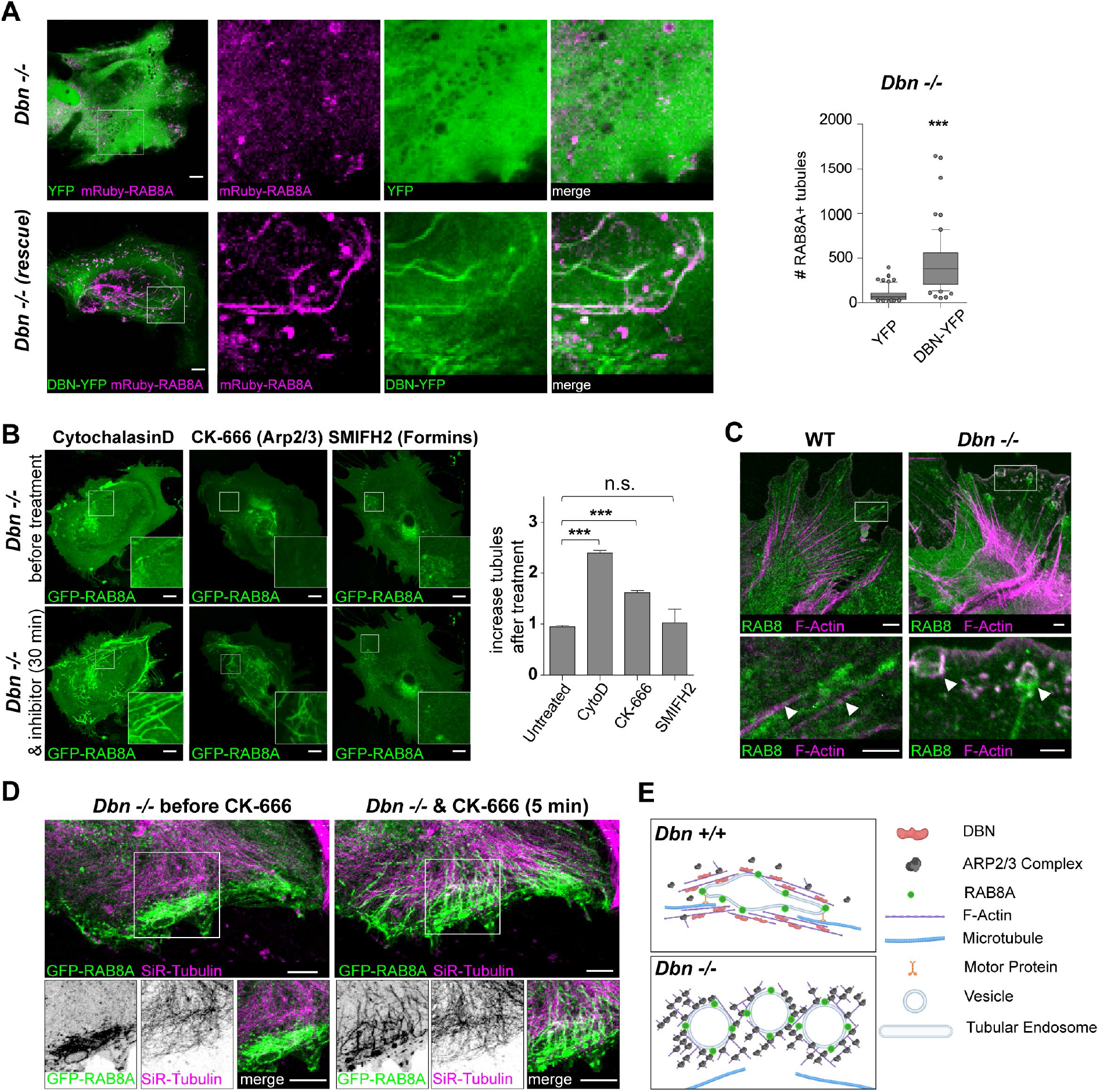
DBN balances the actin nucleation machinery during RAB8A tubule formation. (A) DBN rescues RAB8A membrane tubules in *Dbn*^*−/−*^ astrocytes. *Dbn*^*−/−*^ astrocytes co-transfected with mRuby-RAB8A and YFP-control (upper panels), or with mRuby-RAB8A and DBN-YFP (bottom panels). Scale bars: 10 μm. Bar diagram shows quantification of mRuby-RAB8A tubules; n=63-65 cells from 3 experiments, ***P<0.0001 (Student’s unpaired t-test). (B) Live imaging of *Dbn*^*−/−*^ astrocytes expressing GFP-RAB8A before (upper panels) and 30 min after treatment with different inhibitors (lower panel); 100 nm Cytochalasin D (left), 100 μm CK-666 (center) or 25 μm SMIFH2 (right). Scale bars: 10 μm. Bar diagram shows the quantification of GFP-RAB8A tubules 30 min after inhibitor treatment, Cytochalasin D, n=15-16; SMIFH2, n= 8-9; CK666, n= 9 cells; all from 3 experiments, \*\*\**P*<0.001 (One way ANOVA). (C) Distribution of endogenous RAB8A (green) and F-Actin (magenta) in WT (left panel) and *Dbn*^*−/−*^ (right panel) astrocytes. Arrow heads in WT panel indicate RAB8+ tubules aligned with actin fibers. In the *Dbn*^*−/−*^ panel, arrow heads highlight RAB8+ membrane cisterns. Scale bars: 10 μm. (D) *Dbn*^*−/−*^ astrocytes expressing GFP-RAB8A and labeled with SiR-tubulin before (left panel) and 5 min after adding 100 μM CK-666 (right panel); see Movie 6. Scale bars: 10 μm. RAB8+ tubules formed immediately after adding CK-666 and aligned with adjacent microtubules, while extending into the cytosol. (E) Proposed mechanism. DBN functions as essential switch in the actin network homeostasis, which supports the formation of RAB8A+ tubular endosome along microtubules

### DBN loss affects uptake and intracellular distribution of plasma membranes

RAB8 compartments have previously been reported to engage in plasma membrane recycling as well as in Golgi-derived delivery of cargo to the plasma membrane (Peranen, 2011). To characterize the specificity of endosomal trafficking of the DBN-dependent RAB8 compartment, we analyzed endogenous RAB8-tubules in WT and *Dbn*^*−/−*^ astrocytes after injury using immunocytochemistry. RAB8-positive tubular structures were clearly orientated and extended through the cytosol of WT astrocytes, aligning closely with prominent actin filaments (Figure 4C). In contrast, anti-RAB8 labelling in *Dbn*^*−/−*^ astrocytes produced a diffuse cytosolic signal, with the appearance of cistern-like structures in the cellular periphery. These structures possessed a prominent ring of filamentous actin adjacent to the RAB8 signal, from where stub-like tubules originated (Figure 4C). The presence of peripheral membrane cisterns suggested a general ability of *Dbn*^*−/−*^ astrocytes to endocytose. We provoked increased membrane uptake in GFP-RAB8A expressing astrocytes via EGF-induced macropinocytosis (Wang et al., 2014). EGF-treated WT astrocytes presented throughout the cytosol prominent GFP-RAB8A+ tubules, which locally assembled via the gathering of small Rab8A-positive particles (Movies 4 and 5). In *Dbn*^*−/−*^ astrocytes, RAB8A+ membranes accumulated beneath the plasma membrane, where membrane cisterns frequently emerged but also rapidly disappeared (Figures S6A, Movies 4 and 5). These results establish the function of DBN during the generation and distribution of RAB8 associated membrane tubules,

Given that stabilization of membrane tubules requires the microtubule cytoskeleton in other cell types (Delevoye et al., 2016; Delevoye et al., 2014; Tabdanov et al., 2018), we considered whether excessive ARP2/3 dependent actin dynamics in *Dbn*^*−/−*^ astrocytes (Figure 4B) antagonizes the association of microtubules with the RAB8 compartment. Here, we visualized microtubules using SiR-tubulin in GFP-RAB8A expressing *Dbn*^*−/−*^ astrocytes (Figure 4D, Movie 6). GFP-RAB8A vesicles were visible in the cell’s periphery in proximity to microtubules, but they did show no signs to form tubules. However, during the administration of CK-666, RAB8A-positive vesicles turned immediately into tubules, which then extended along the microtubules towards the cell body (Figure 4D). Thus, DBN controls RAB8 membrane trafficking by antagonizing ARP2/3 dependent actin networks to enable the transport of tubular membranes along microtubules. In turn, DBN-loss disturbs the normal actin equilibrium and creates excessive ARP2/3 activity, which prevents the microtubule-assisted formation of tubular endosomes from RAB8+ vesicles (Figure 4 E).

### DBN-dependent RAB8A tubules are required for β1-integrin trafficking

A known RAB8 target in cell lines is β1-integrin, a key molecule in cell polarization and migration (Hattula et al., 2006; Sun et al., 2016), which also controls astrocyte differentiation and reactivity (Hara et al., 2017; North et al., 2015; Robel et al., 2009). We used a C-terminus-specific β1-integrin antiserum to detect the spatiotemporal distribution of β1-integrin in astrocytes during injury, and identified prominent β1-integrin foci in RAB8 membrane tubules (Figure 5A). To examine if DBN regulates β1-integrin, as a first read-out, we exploited the composition and assembly of focal adhesions, which are hot spots of β1-integrin activity. Antibody labeling in WT astrocytes during scratch injury showed the concentration of active β1-integrin, as well as the intracellular adapter paxillin in mature focal adhesions (Sun et al., 2016). *Dbn*^*−/−*^ astrocytes, in contrast, exhibited scattered membrane distribution of active β1-integrin with smaller paxillin+ focal adhesions (Figure 5B). By tracking of GFP-tagged paxillin, we identified a persisting reduction in focal adhesion sizes during the injury-induced polarization of *Dbn*^*−/−*^ astrocytes, when compared to WT cells (Figure 5C). Thus, DBN controls the presentation of β1-integrin to focal adhesions in astrocytes during their responses to injury.

**Figure 5:**
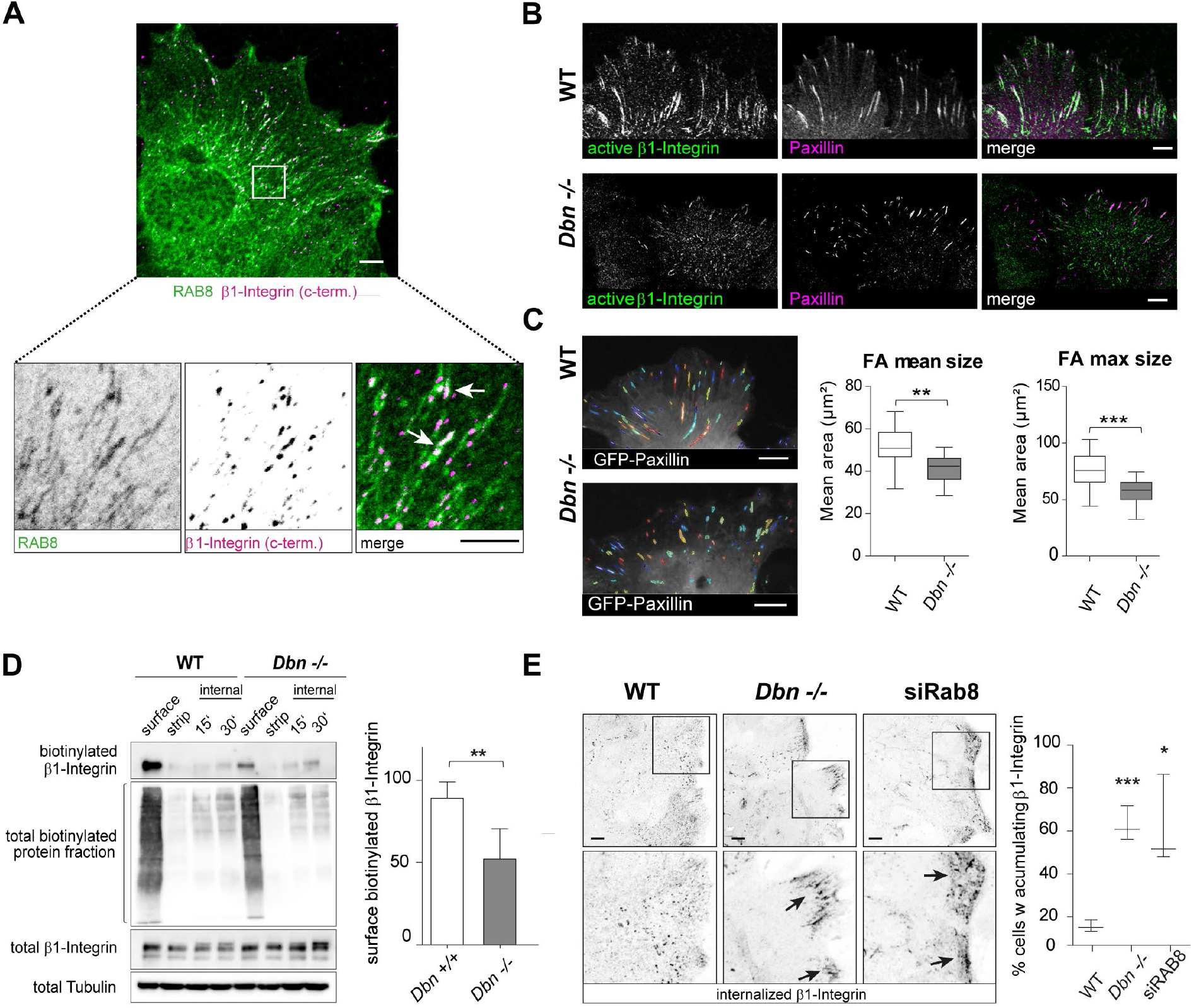
DBN regulates RAB8 membrane trafficking of β1-integrin during injury. (A) Detection of β1-integrin (magenta in merged images) in RAB8+ tubules (green in merged images) in astrocytes. Scale bar: 10 μm. (B) Confocal images of WT (upper panel) and *Dbn*^*−/−*^ a*s*trocytes (bottom panel) after mechanical injury, labeled for active β1-integrin (green) and paxillin (magenta). Scale bars: 10 μm. (C) Focal adhesions in WT (upper panel) and *Dbn*^*−/−*^ astrocytes (lower panel) expressing GFP-paxillin analyzed by the Focal Adhesion Analysis Server. Bar diagram shows quantification of focal adhesion (FA) mean sizes and FA maximum sizes over 22 hours during live imaging experiments (n= 17 cells from 3 independent experiments, \*\**P*<0.01, *** *P*<0.001). (D) Streptavidin-pulldown of proteins after surface biotinlyation of injured WT and *Dbn*^*−/−*^ astrocytes. Western blot shows biotinylated β1-integrin in different conditions: immediately after surface biotinylation (surface), after removal of the surface-biotin label with MESNA (strip), 15 min and 30 min after incubating labeled astrocytes at 37°C followed by MESNA treatment to remove the fraction of surface-exposed labeled proteins. Bar diagram shows quantification of surface β1-integrin after normalization to total integrin levels; n= 6 experiments \*\**P*<0.01, (Student’s unpaired t-test). (E) Labeling of internalized ligand-bound β1-integrin following antibody feeding in WT astrocytes (left image), *Dbn*^*−/−*^ astrocytes (center image) and WT astrocytes after siRNA depletion of RAB8A and RAB8b (right image). Bar diagram shows quantification of astrocytes with antibody-labeled β1-integrin at leading edges; boxes in upper panels are magnified in lower panels. n=20-37 cells from 3 experiments, * *P*<0.05, *** *P*<0.001 unpaired Student’s test).

We then investigated if the DBN-dependent, injury-induced RAB8 membrane tubules are required for β1-integrin trafficking. To study the distribution and trafficking of β1-integrin independent of conformational states and antibody epitopes, we performed surface biotinylation experiments in injured WT and *Dbn*^*−/−*^ astrocyte cultures. Whilst signals of intracellular biotinylated β1-integrin were at the detection limit after internalization, we found a substantial reduction of surface β1-integrin in *Dbn*^*−/−*^ astrocytes when compared to WT astrocytes (Figure 5D). Finally, we followed the internalization of active integrin *in situ* with an antibody-feeding assay. Thirty minutes after antibody labeling with anti-β1-integrin, the receptor was widely distributed throughout WT astrocytes (Figure 5E). In contrast, in *Dbn*^*−/−*^ astrocytes, β1-integrin accumulated beneath the plasma membrane of polarizing cells. We obtained a comparable β1-integrin distribution in WT astrocytes after depleting both RAB8 isoforms with RNAi (Figures 5E), suggesting that DBN-dependent RAB8 compartments are major routes for β1-integrin trafficking. We conclude that the DBN-dependent, injury-induced RAB8 tubule compartment functions as an important hub to distribute internalized β1-integrin in astrocytes during injury.

### DBN loss induces intracellular membrane accumulation in reactive astrocytes *in vivo*

The extensive accumulation of RAB8+ membranes beneath the leading edge of cultured D*bn*^−/−^ astrocytes prompted us to analyze internal membranes *in vivo* at the ultrastructural level using high-resolution transmission electron microscopy (TEM). We compared WT and *Dbn*^*−/−*^ brains from 7-days post stab-injury, using GFAP+ to locate reactive astrocytes in lesions. Processes of reactive astrocytes in WT brains in proximity to the stab lesion contained several endosome-like organelles, which frequently connected to thin membrane tubules. In contrast, processes of *Dbn*^*−/−*^ reactive astrocytes in proximity to the stab-injury displayed massive cytoplasmic vacuolization. The vacuoles resembled multilamellar bodies, as they were filled with amorphous, electron-dense material that was surrounded by double or multiple concentric membranes layers (Figure 6A) (Hariri et al., 2000). Quantification of multilamellar bodies at injury and contralateral sites of WT and *Dbn*^*−/−*^ brains showed that this compartment was induced upon injury only in the absence of DBN (Figures 6A, B). In addition, we investigated the ultrastructure of astrocytic endfeet, which sustain high levels of membrane trafficking during nutrient and metabolite uptake, and efflux of waste products (Schiweck et al., 2018). Analogous to astrocyte processes, astrocyte endfeet at the injury site in WT brains contained several endosomal structures. In contrast, excessive amounts of multilamellar bodies filled the endfeet at injury sites in *Dbn*^*−/−*^ mice (Figure S6B). In summary, astrocytes from *Dbn*^*−/−*^ mice show excessive, injury-specific accumulation of membrane-derived material in both astrocytic processes and endfeet.

**Figure 6:**
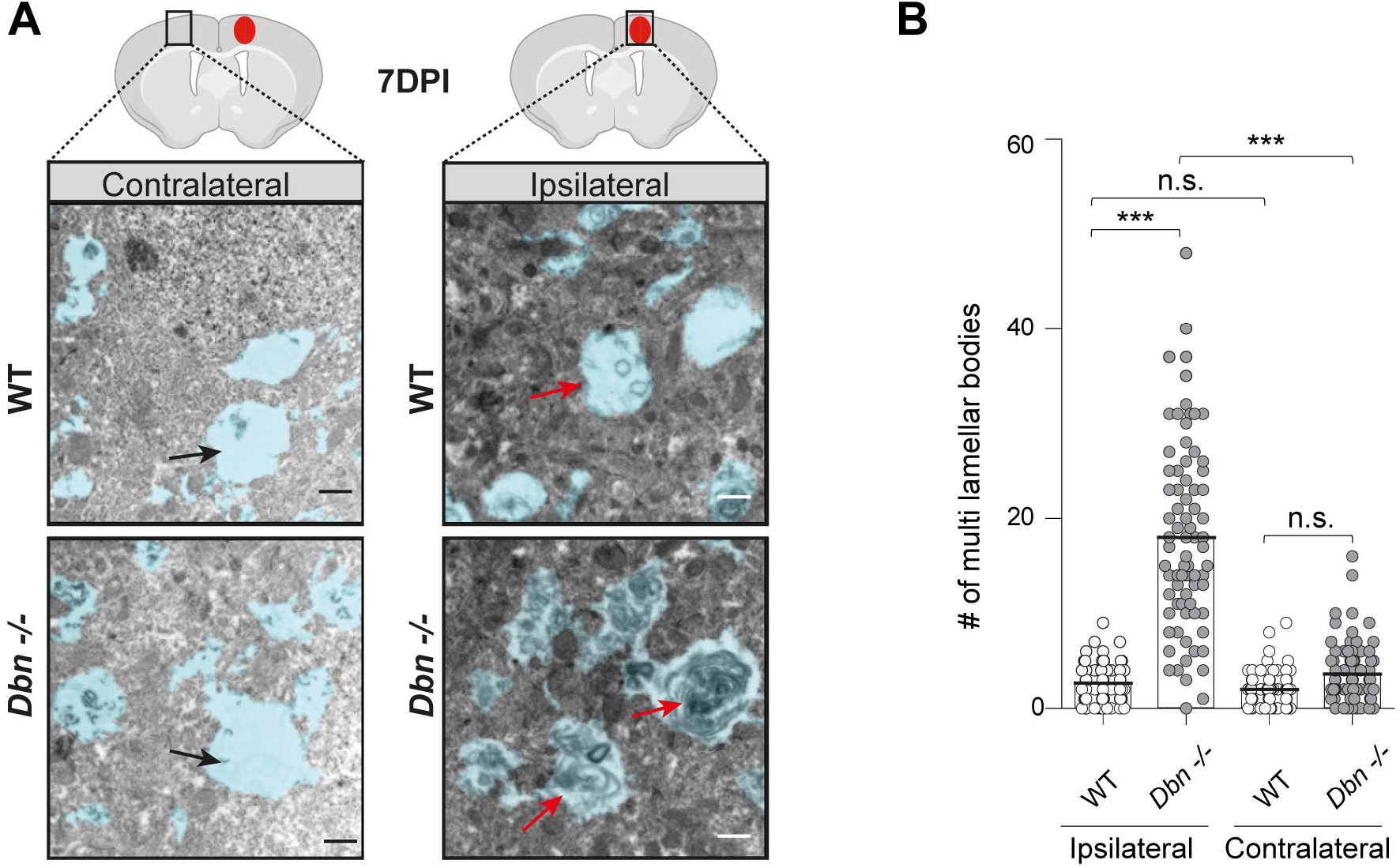
DBN loss induced membrane accumulations in polarizing astrocytes during injury. (A) TEM ultrastructural architecture of astrocytes in brains of WT and *Dbn*^*−/−*^ mice. Images were acquired on the contralateral side of injury (left panels) or inside the lesion core (right panels); 7 days post stab-injury. Astrocyte processes are shaded in blue. Scale bars: 500 nm. (B) Quantification of multilamellar bodies. Field of view size: 1170 μm^2^. Number of fields/view: 92 WT and 77 *Dbn*^*−/−*^ ipsilateral, 85 WT and 77 *Dbn*^*−/−*^ contralateral from 3 WT and 3 *Dbn*^*−/−*^ animals. \*\*\**P*<0.001 (ANOVA).

## Discussion

We have identified a novel function of DBN in protecting the brain from tissue damage following injury. We show that (1) under physiological conditions, DBN protein is not expressed in astrocytes, however its injury-induced upregulation in reactive astrocytes is required for the coordinated formation and maintenance of glial scars, demonstrating its essential role in effective tissue protection in the brain. (2) At the cellular level, DBN shifts the actin network organization from ARP2/3 dependent arrays to microtubule-compatible scaffolds, which facilitate the formation of injury-induced RAB8A-positive membrane tubules. (3) These tubules serve as a hub for the membrane trafficking of surface proteins involved in coordinating adhesive responses, such as β1-integrin. (4) By the actin-dependent facilitation of RAB8 membrane tubules, DBN is essential for the maintenance of astrocyte reactivity and prevention of neurodegeneration by ensuring the physical integrity of the glial scar. We therefore propose a conceptually new role for DBN as an injury-induced actin regulator in reactive astrocytes.

### DBN functions as an injury-induced actin regulator in membrane trafficking

DBN has been identified as highly abundant in dendritic spines and developing neurites, and confers resilience in dendritic spines during cellular stress responses (Dun et al., 2012; Koganezawa et al., 2017; Kreis et al., 2019). Unexpectedly, the cell biological mechanism underlying the role of DBN during glial scar formation involves a directive function for its participation in membrane trafficking. Until now, few studies have linked DBN to membrane dynamics in cells, other than a demonstration that DBN principally binds to membranes (Xu and Stamnes, 2006), that it restricts the entry of rotavirus into cells by limiting endocytosis (Li et al., 2017), and that it regulates antigen presentation of dendritic cells (Elizondo et al., 2019). Our study identifies a further avenue in DBN-dependent membrane trafficking and protein sorting mechanism that functions by stabilizing RAB8-positive tubular endosomes upon injury.

In cell culture and *in vivo*, we discovered prominent RAB8-positive tubular endosomes specific to a post-injury setting in astrocytes, which extend long palisading-like processes into wound areas. The GTPases RAB8A and RAB8B facilitate distinct routes of polarized membrane transport. RAB8 is involved in endocytosis, membrane recycling, autophagy and exocytosis by associating with vesicles, macropinosomes and tubules (Peranen, 2011) (Grigoriev et al., 2011; Ryan and Tumbarello, 2018). Moreover, RAB8 has been implicated in various diseases ranging from microvillar inclusion disease (MVID) and cancer to neurodegenerative diseases such as Alzheimer’s and Parkinson disease (Peranen, 2011). Tubular membrane compartments serve as logistical platforms to sort membrane-bound cargos (van Weering and Cullen, 2014). Membrane tubulation occurs particularly on maturing early endosomes and macropinosomes to separate components from the quick ‘bulk flow’ back to the plasma membrane and to direct them towards other compartments such as the Golgi network or recycling endosomes (Naslavsky and Caplan, 2018). The actin cytoskeleton creates stable tubular subdomains, where membrane-bound cargos segregate and concentrate according to their designated destinations (Bowman et al., 2016; Puthenveedu et al., 2010; Simonetti and Cullen, 2019). We propose a role for DBN in inward-directed RAB8-based membrane trafficking: upon wounding, DBN may contribute to the uptake of plasma membrane material by compiling endocytosed receptors and material in the injury-induced RAB8-compartment that are essentially sorted by DBN-stabilized tubules (Figure 7).

**Figure 7:**
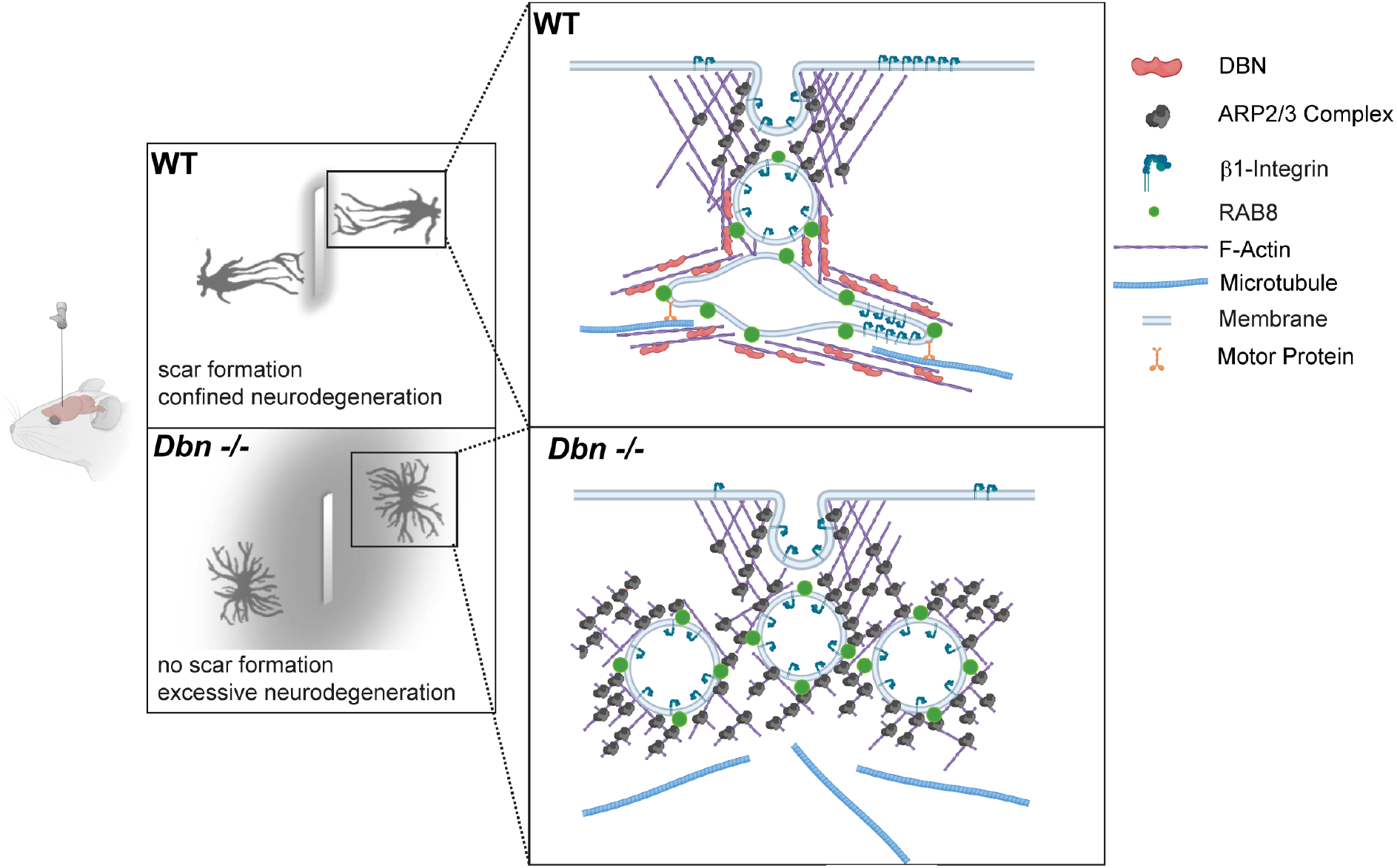
Astrocytes require DBN to control brain damage during brain injury. Proposed model of how DBN/Rab8 functions during astrocyte polarization and scarring responses. In response to CNS injury, DBN generates actin scaffolds suitable for tubule-based membrane trafficking. DBN-loss results in the accumulation of endocytotic vesicles beneath the plasma membrane and affects sorting of adhesion receptors essential for interaction with lesion core components. In this way, DBN/Rab8 membrane tubules are fundamental for the polarization and scarring responses of reactive astrocytes.

### DBN antagonizes ARP2/3-dependent actin dynamics

ARP2/3-dependent actin nucleation has been shown to drive the fission of existing tubules into vesicular endosomes by the WASH complex (Derivery et al., 2009). DBN associates dynamically with scaffolds of forming tubular endosomes and counteracts their ARP2/3 dependent fission (Figure 4B,D). DBN could thereby occlude ARP2/3 binding sites on the actin scaffold by its sidewise binding to actin filaments (Grintsevich et al., 2010). This model is consistent with DBN competition of ARP2/3 function, as identified in our pharmacological rescue experiments. In addition, DBN could also facilitate the extension of RAB8 tubules from underneath the leading edge into the cell body by arranging the surrounding actin architecture. The observable actin filaments in WT astrocyte processes generally run parallel to RAB8 tubules. These linear actin filaments may serve as permissive scaffold for RAB8 tubules, which then stabilize along intermingled microtubules analogous to membrane tubulation during sorting to recycling endosome transition in non-neuronal cells (Delevoye et al., 2016; Delevoye et al., 2014; Tabdanov et al., 2018). This mechanism would be susceptible to mislocalized ARP2/3-actin arrays upon DBN loss. Our live imaging results also support this mechanism in view of *Dbn*^*−/−*^ astrocytes, where, until their pharmacological disruption, ARP2/3-actin networks prevent the tubulation and transport of RAB8A-positive membrane along microtubules. We propose that DBN serves as master switch to enable the assembly of microtubule-compatible actin networks for tubular membrane trafficking (Figure 7).

### Cellular trafficking by RAB8 tubular endosomes

Tubular endosomes and/or tubule-derived vesicles deliver their content to intracellular destinations, where they may be redistributed or degraded according to cellular requirements for efficient polarized astrocyte outgrowth. DBN deficiency disrupts the RAB8 dependent tubular protein sorting machinery. Consequently, membranes and associated proteins accumulate in intermediate endosomal compartments instead of being rerouted in a polarized manner in sufficient numbers. The resulting deficits and mislocalization of surface receptors first evokes the failed wounding response and polarization of astrocytes followed by their erratic behavior during scarring. The disarray within the scarring astrocytes could lead to the downregulation of their reactivity, as shown by the loss of GFAP– either in a cell-autonomous manner and/or by an affected interplay with other cell types such as microglia.

Our protein traffic-based mechanism is supported by findings in DBN-depleted epithelial cells, which share some localization defects in apical markers with RAB8A-deficient intestinal cells during microvillar inclusion disease (Sato et al., 2007; Vacca et al., 2014). In addition, RAB8 was shown to organize cell adhesion in conjunction with RAB13 by transporting adhesion molecules, in cultured epithelial cells (Sakane et al., 2012). Moreover, the closely RAB8-related RAB13 organizes the collective cell migration of these non-neuronal cells in wounding assays in an ordered manner (Sakane et al., 2016), analogous to our findings with erratic DBN-deficient astrocytes in glial scars *in vivo* and during live imaging experiments.

### β1-Integrin trafficking and astrocyte reactivity

The induction of tubular endosomes in response to injury suggests that they represent an efficient way of transporting and sorting cargo relevance for the injury response in astrocytes. We show the mislocalization of endogenous β1-integrin in DBN-deficient astrocytes in a Rab8-dependent manner. Cells require tight control of β1-integrin in terms of levels and distribution at the cell surface to migrate efficiently. The key mechanism for this is retrograde trafficking (Shafaq-Zadah et al., 2016). Inadequate β1-integrin subunits at the surface impair the assembly and turnover of focal adhesions, as seen with the outgrowth defects observed in DBN-deficient astrocytes. In particular, the vanishing astrocyte reactivity highlights comprehensive defects in signal reception and computation through the mis-sorting of membrane receptors. Astrocytic β1-integrin plays an important role as a co-receptor in signaling pathways, which control astrocyte reactivity (Hara et al., 2017; North et al., 2015; Robel et al., 2009). However, other mis-regulated membrane receptors undoubtedly also contribute to the observed phenotype, and further research screening DBN-deficient astrocytes is required to identify them.

### DBN function in brain astrocytes

A major finding of this study is the relevance of injury-induced DBN in forming functional glial scars. We showed that DBN is crucial for the coordinated polarization and palisade-like outgrowth of astrocytes and ultimately, the formation of glial scars. The establishment in turn, of a glial scar, is absolutely essential for damage containment in the CNS (Wanner et al., 2013), as highlighted by the high neurodegeneration observed in DBN-deficient mice without glial scars. What is more, the severity of the DBN-dependent membrane trafficking defects became apparent in our *in vivo* TEM studies. We showed the prominent enrichment of membranous material in multi-lamellar structures within processes and endfeet of DBN-negative astrocytes. The extent of these accumulating multilamellar structures in astrocytes is, to our knowledge, unique, occurs only upon injury and supports our results of trafficking defects in cell culture. Moreover, this phenomenon resembles membrane whorls, which are evocative for autophagy and accumulate in smooth muscle cells, in particular during atherosclerosis (Hariri et al., 2000; Martinet and De Meyer, 2009). Analogous structures have been reported from Alzheimer’s disease and amyotrophic lateral sclerosis patients as well as neurons expressing mutated huntingtin (Kegel et al., 2000; Nixon et al., 2005; Sasaki, 2011). Our findings indicate a putative role of the polarized DBN-Rab8-dependent membrane trafficking for disease-specific autophagy, which is well aligned with astrocytes becoming phagocytic under pathological conditions (Morizawa et al., 2017). The aspects of DBN-deficiency identified in astrocytes in our study, are likely to exacerbate a mild CNS insult into major trauma with progressing neurodegeneration. Further dissection of the underlying mechanism could open new therapeutic avenues to treat CNS trauma and, possibly, degenerative conditions, which to date have dismal outcomes.

In conclusion, this study identifies an important function of injury-induced DBN in controlling damage containment in the CNS via membrane trafficking. Since DBN and its downstream targets RAB8 and β1-integrin are broadly expressed, it is conceivable that this mechanism may be pivotal in other pathologies in the CNS as well as in different organ systems.

## Acknowledgments

We would like to thank Magdalena Götz and Sofia Grade (LMU Munich) for provision of brain slices, expert opinion and critical feedback on the manuscript. We thank Heike Heilmann, Kerstin Schlawe, Kristin Lehmann and Beate Diemar for excellent technical assistance. We would like to thank the Advanced Medical BioImaging Core Facility (AMBIO) and the NeuroCure multi-user Microscopy Core Facility for usage of microscopes. We thank Dietmar Schmitz for providing the B6.Cg-Tg(Camk2a-cre)T29-1Stl/J mouse line. Funding was provided by the DFG SFB 958, A16 (B.J.E.); SFB TRR 186, A10 (B.J.E.); NeuroCure EXC257 (B.J.E.); Sonnenfeld Stiftung (J.S.).

## Author Contributions

Conceptualization, B.J.E and K.M.; Investigation, J.S., K.M., A.M.W. and J.L.; Methodology, K.M. & J.S.; Software, J.S.; Formal analysis: J.S. and K.M.; Writing and Editing, B.J.E., K.M. and J.S.; Visualization, K.M., J.S; Funding Acquisition, B.J.E.; Supervision, B.J.E., K.M. and I.V.

## KEY RESOURCES TABLE

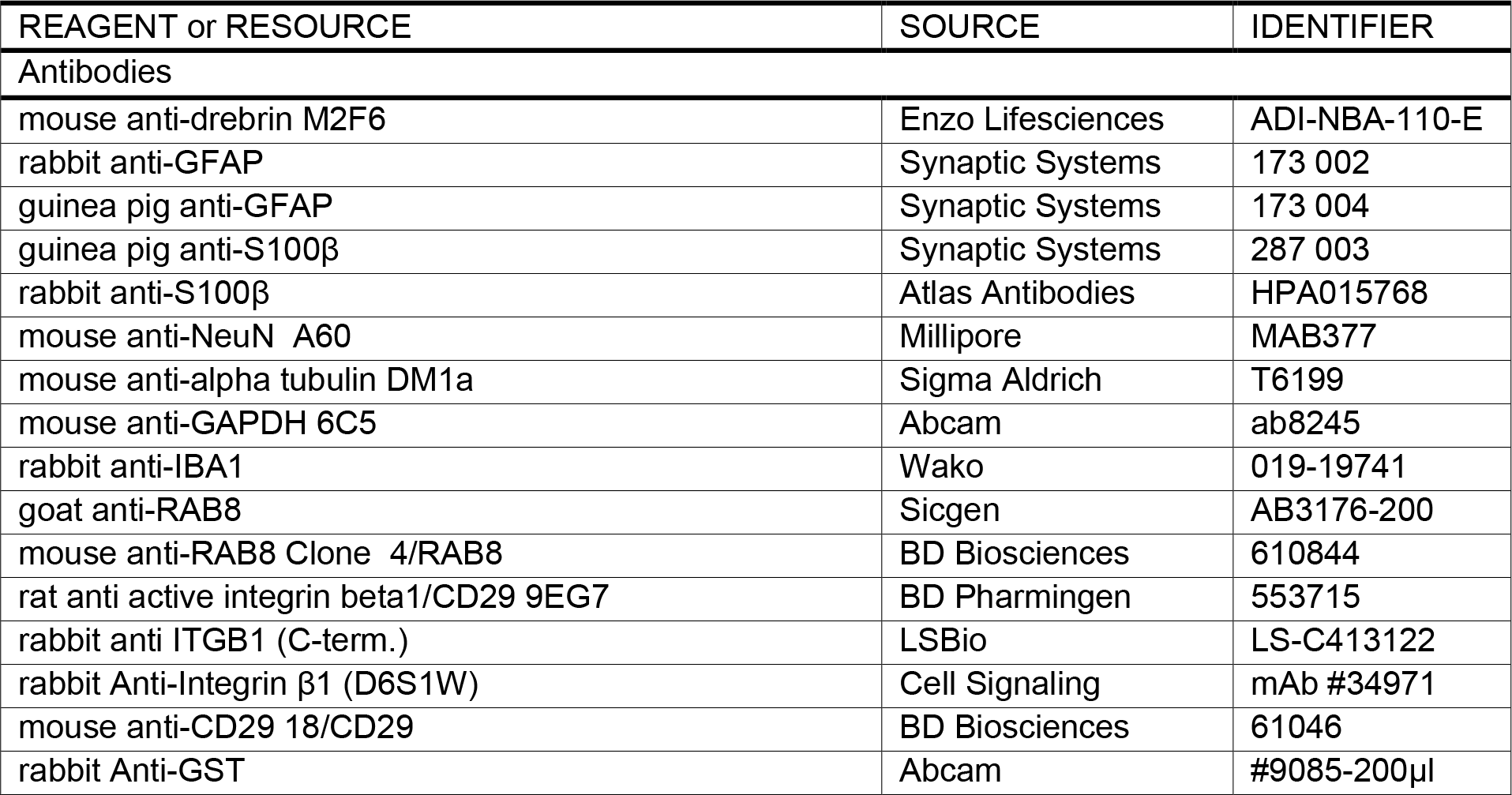
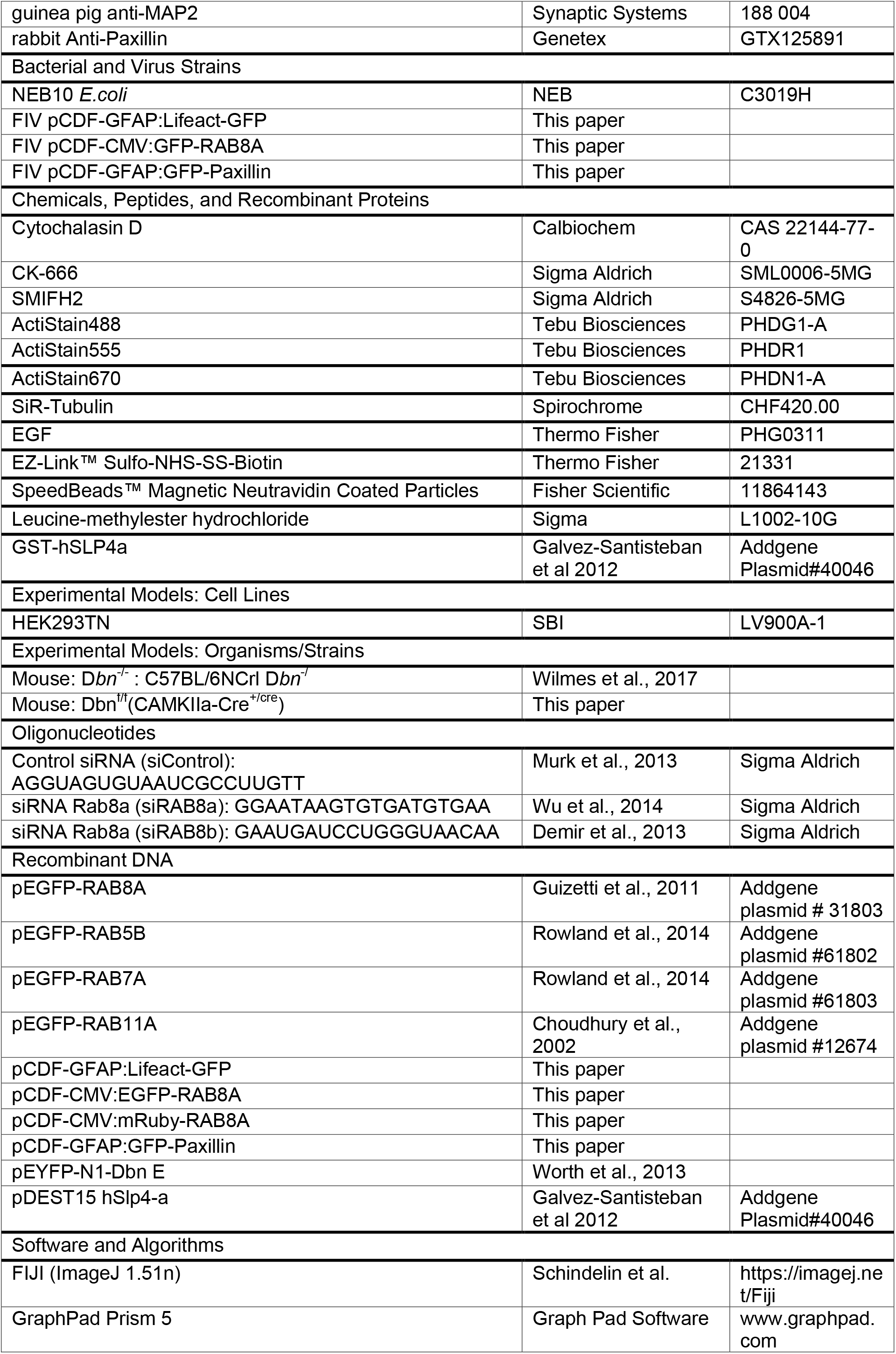
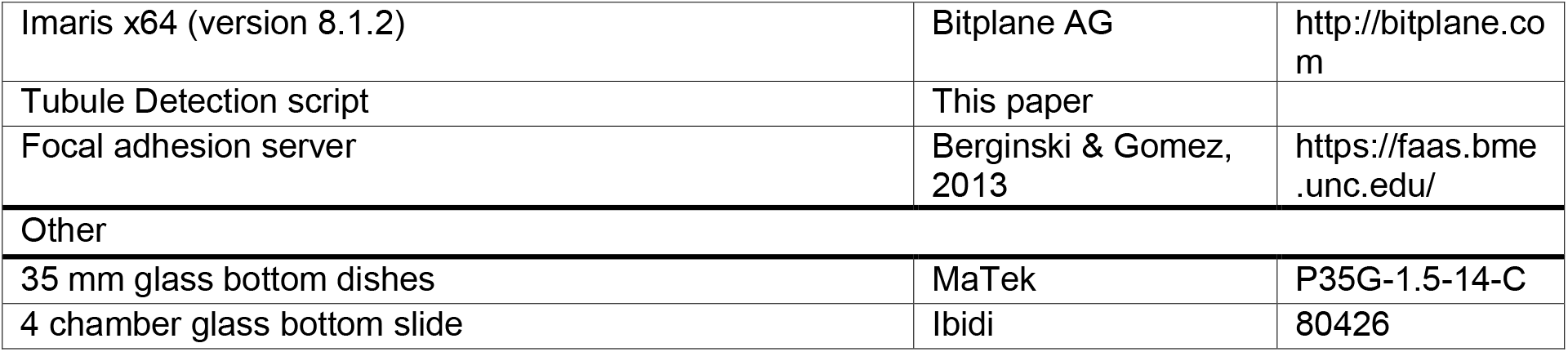

## Experimental Procedures

### Ethical Approval

All animals were handled in accordance with the relevant national guidelines and regulations. Protocols were approved by the ‘Landesamt für Gesundheit und Soziales’ (LaGeSo; Regional Office for Health and Social Affairs) in Berlin, and animals are under the permit number G0189/14.

### Mouse Strains

*Dbn*^*−/−*^ mice were generated and described previously (Willmes et al., 2017). B6.CAMK:Cre/Dbn^fl/fl^ mice were generated by crossing B6.Cg-Tg(Camk2a-cre)T29-1Stl/J (kindly provided by Dietmar Schmitz, Charité Berlin; (Tsien et al., 1996) with B6. Dbn^fl/fl^ mice.

### Antibodies & Reagents

Mouse anti-DBN M2F6 (Enzo Lifesciences, ADI-NBA-110-E, IF & IHC 1:100; WB 1:1000), rabbit anti-GFAP (Synaptic Systems, 173 002, IF & IHC 1:1000), guinea pig anti-GFAP (Synaptic Systems, 173 004, IHC 1:400), guinea pig anti-S100β (Synaptic Systems, 287 003, IHC 1:200); rabbit anti-S100β (Atlas Antibodies, HPA015768, IHC 1:1000), mouse anti-alpha tubulin DM1a (Sigma Aldrich, T6199, WB 1:5000), mouse anti-NeuN A60 (Millipore, MAB377, IHC 1:1000), rabbit anti-IBA1 (Wako, 019-19741, IHC 1:500), anti-MAP2 (Synaptic systems, 188 004, IHC 1:500), mouse anti-GAPDH 6C5 (Abcam, ab8245, Wb 1:5000), mouse anti-RAB8 (BD Biosciences, 610844, WB 1:1000), goat anti-pan RAB8 (Sicgen, AB3176-200, IF & IHC 1:100), rabbit anti-integrin beta1 (Cell Signaling, #4706S, WB 1:1000), mouse anti-CD29 18/CD29 (BD Biosciences, 61046, WB 1:1000), rat anti-active integrin beta1/CD29 9EG7 (BD Pharmingen, 553715, IF 1:100), rabbit anti-integrin beta1 c-term (LSBio, LS-C413122, IF 1:100), rabbit anti-paxillin (Genetex, GTX125891, IF (1:250), rabbit anti-GST (Abcam, #9085-200μl, WB 1:500). Horseradish peroxidase (HRP)-conjugated secondary antibodies were purchased from Millipore. Cross-absorbed secondary antibodies conjugated to cyanine or Alexa dyes were purchased from Dianova and Thermo Scientific, respectively. Filamentous actin was labeled with ActiStain488, 555 or 670 phalloidin (Tebu Bio). DNA staining was carried out using Hoechst 33258 (Thermo Scientific).

### Plasmids

Lifeact-GFP was cloned into the pCDF backbone (SBI), after replacing copGFP with a suitable multiple cloning site via XbaI/SalI digest (Sequence of MCS: TCTAGAGCTAGCGCTACCGGTCGCCACCATGGGATGTACAGCGGCCGCGTCGAC). Accordingly, the CMV promoter was substituted with the astrocyte-specific gfaABC1 promoter, kindly provided by Michael Brenner (University of Alabama, US), using PCR (pGFAP-5:ATAGATATCAACATATCCTGGTGTGGAGTAGGG, pGFAP-3:ATAGCTAGCGCGAGCAGCGGAGGTGATGCGTC). pEGFP-RAB8A was a gift from Daniel Gerlich (Addgene plasmid # 31803, (Guizetti et al., 2011). EGFP-Rab8a was cloned via PCR into the pCDF backbone for viral expression (pCDF-GFP-5:GACCTCCATAGAAGATTCTAGAGCTAGCATGGTGAGCAAGGGCGAGGAGCTGTTC, pCDF-RAB8A-3: GTAATCCAGAGGTTGATTGTCGACTCACAGAAGAACACATCGGAAAAAGCTGC). mRuby-RAB8A was generated through replacement of EGFP in pCDF-eGFP-RAB8A with mRuby via NheI/BsrgI from pcDNA3-mRuby kindly provided by Nils Rademacher (Charité Universitätsmedizin Berlin, Germany). The lentiviral construct pCDF-paxillin-EGFP was subcloned from pEGFP-N3-paxillin (gift from Rick Horwitz, (Laukaitis et al., 2001); Addgene plasmid #15233) into pCDF-GFAP:Lifeact-GFP via NheI/BsRGI after silent mutation of an internal BsRGI site (Paxillin-BsrGImut-s: GAGGAGGAACACGTGTATAGCTTCCCAAACAAGCAG; Paxillin-BsRGImut-as: CTGCTTGTTTGGGAAGCTATACACGTGTTCCTCCTC).

Previously published pEYFP-N1-DBN E was co-transfected in rescue experiments in *Dbn*^*−/−*^ astrocytes in conjunction with pCDF-mRuby-RAB8A (Worth et al., 2013). pEYFP-N1 (Takara Bio Inc) was used as corresponding negative control.pDEST15-hSlp4-a was purchased from Addgene (Plasmid#40046).

### Cell Culture & Plasmid Transfection

Cortical astrocytes were isolated from mouse brains according to procedures for rats (Murk et al., 2013) and plated on plastic or glass-bottom dishes (MaTek Corporation or ibidi) coated with 0.025% collagen type I (BD Biosciences) and 100 μg/ml poly-ornithine (Sigma-Aldrich). Depletion of microglia via 60 mM leucine methylester (LME) in complete medium was carried out for 60-90 min. Enriched microglia cultures were acquired by shaking T75 cm flasks with polygonal astrocytes at 150 rpm at 37°C for 2 h prior to treatment with LME. Supernatant with floating microglia were plated on pLL-coated glass coverslips. Microglia cultures contained less than 5% astrocytes. Mixed neuronal cultures from mouse cortices were prepared and cultured as published previously (Schrotter et al., 2016). HEK293TN were obtained from BioCAT (Cat. No. LV900A-1-GVO-SBI). All culturing materials were sterile and cell culture techniques were undertaken in Class II vertical laminar flow cabinets (ThermoFisher Scientific). HEK293TN cells were not authenticated. Cultured cells were not tested for mycoplasma. Astrocytes were transfected using TransIT LT1 (Mirus) according to the manufacturer’s protocol.

### Lentivirus production and astrocyte infection

Probes of fluorescent RAB8A and paxillin were expressed in astrocytes via lentiviral transduction. FIV particles were produced in the HEK293TN producer cell line (BioCat) and harvested analogously to HIV particles (Wittenmayer et al., 2009). 70% confluent astrocytes from WT and *Dbn*^*−/−*^ mice on glass bottom dishes were transduced seven days before scratch wounding and live cell imaging.

### RNA interference

Astrocytes were transfected with siRNAs using Lipofectamine RNAiMax (Thermo Scientific) according to the manufacturer’s instructions. The following published siRNA oligonucleotides were used (Sigma-Aldrich): siRab8a (GGAATAAGTGTGATGTGAA (Wu et al., 2014), siRab8b (GAAUGAUCCUGGGUAACAA (Demir et al., 2013), and control siRNA (AGGUAGUGUAAUCGCCUUGTT, (Murk et al., 2013).

### Scratch wound *in vitro* and live cell imaging

Confluent astrocytes were cultured in phenol red-free DMEM (Thermo Scientific) with 10% FBS. Scratch wounds were introduced as described previously (Etienne-Manneville and Hall, 2001). Injured astrocytes were either biochemically analyzed via Western blot or studied in live cell imaging. Cell behavior was followed by live cell imaging for 22 or 30 h using a Nikon Widefield with CCD camera, scanning stage and environmental control chamber (OKO lab) and a 40x objective (N.A. 0.7x) with 1.5x intermediate magnification. Cells were imaged at 20-min intervals with composite large images (3×3 fields of view). Imaging start, unless specified otherwise, was 4 h after *in vitro* injury. Astrocytes in compiled movies were analyzed using kymographs in Fiji as described previously (Hiroyasu et al., 2016).

### Quantitation of membrane tubules

Tubular membrane compartments were quantified based on a macro for Fiji, originally designed to analyze analogous structures in still images in heart muscle (Pasqualin et al., 2015). We extended the functionality of the original code by enabling analyses of z-stack image sequences acquired in live imaging experiments. The script is provided in the Supplementary Data.

### Pharmacological treatments

100 nM Cytochalasin D (Merck Calbiochem, Cat. No. 250255), 100 μM CK-666 (Sigma-Aldrich, Cat. No: SML0006-5MG) or 25 μM SMIFH2 (Sigma-Aldrich, Cat. No. S4826-5MG) were pre-diluted in phenolred-free DMEM and added to GFP-RAB8A-expressing astrocytes after recording baseline for 30 min without treatment. Microtubule dynamics were imaged after incubating astrocytes for six hours in 1 μM SiR-tubulin (Spirochrome, cat#: SC002) in phenolred-free DMEM.

### Immunocytochemistry

Cultured astrocytes were fixed using 3.7% formaldehyde in cytoskeleton-preservation buffer (25 mM HEPES, 60 mM PIPES, 10 mM EGTA, 2 mM MgCl_2_, pH 7.4) for 20 min. After three washes with cytoskeleton-preservation buffer, cell permeabilization, while maintaining the utmost integrity of the cytoskeleton and endosomes, was achieved through 3 min incubation with 0.02% Triton-X100 in PBS. After three washes with PBS and brief incubation in 1% BSA in PBS, cells were incubated with primary antibodies diluted in PBS for 1 h at RT. After three washes with PBS and brief incubation in 1% BSA in PBS, cells were incubated in highly cross-absorbed secondary antibodies for another hour at RT. After three washes with PBS, cells were mounted in Mowiol. To visualize endogenous RAB8 tubules, PBS was generally replaced by cytoskeleton-preservation buffer. Cells were permeabilized by incubation with 0.02% Triton-X100 in cytoskeleton-preservation buffer for 30 min at RT. Wash steps were extended to 5 min each.

### Confocal Microscopy & Imaging Processing

Cells were imaged via multitrack mode on either Leica Sp8 (Leica) or Nikon A1Rsi+ (Nikon) confocal microscopes. Large image and multipoint scans were performed on Nikon A1 microscopes using an automated stage. Image processing were performed in FiJi and/or Imaris (Bitplane). Kymographs were generated and analyzed via FiJi. Microglia morphometry was analyzed based on IBA1 immunoreactivity and the surface algorithm of Imaris. Movies were compiled and annotated with Hitfilm Express 13 (FXhome Limited).

### Focal Adhesion Analysis

To assess focal adhesion size, primary astrocytes were transduced with FIV lentivirus encoding for pCDF-GFAP:paxillin-EGFP. Seven days after transduction, medium was changed to Phenol-free DMEM and scratch injury was performed, followed by live cell imaging for 22 hours in 20 minute intervals.

Acquired images were submitted to the Focal Adhesion Analysis Server (Berginski and Gomez, 2013) with the following specifications: Imaging frequency: 20 min; Min. adhesion size 20, Max. adhesion size 500; all further settings were left at default. The analyzed files obtained from the adhesion server were used to identify a Region of Interest at the leading edge of astrocytes extending into the scratch. Each Region of Interest contained at least 10 focal adhesions. Using the open source software INKSCAPE, the assigned number of every analyzed focal adhesion was identified and matched with the corresponding data (Maximum adhesion size, mean adhesion size) from the excel-sheet as described on the focal adhesion analysis server website (https://faas.bme.unc.edu/results_understanding).

### Surface biotinylation & internalization assay

Cultured astrocytes were incubated in 1 mg/ml cell-impermeant EZ-Link Sulfo-NHS-SS-Biotin (Thermo Fisher) in ice-cold PBS pH 8 for 2 h. Any remaining unreacted biotin was removed by three washes with ice-cold PBS followed by 10 min of quenching with 100 mM glycine in ice-cold PBS. Cells were then split into four groups: (1) directed analyses of surface biotinylated proteins after labeling and quenching, (2) specificity control of surface biotinylation by removing biotin surface labeling with 50 mM 2-mercaptoethanesulfonic acid sodium salt (MESNA) in wash buffer (150 mM NaCl, 0.2% BSA, 20 mM Tris, pH 8.6), (3) pulse-chase analyses of internalized surface proteins after 15 min or (4) 30 min incubation at 37°C in complete medium followed by MESNA-dependent removal of the surface-label. After three washes with wash buffer, samples were treated with 50 mM MESNA in wash buffer for 1 h on ice. Cell lysis of all samples was performed after two washes with ice-cold PBS by incubating cells for 20 min in ice-cold 1% Triton X100 in PBS with protease inhibitor cocktails (Merck, Calbiochem set III, Cat. No. 539134), subsequent thorough scraping and sonication.

Biotinylated proteins were isolated by incubating the samples with equal total protein amounts overnight at 4°C and gentle agitation with 30 μl of Neutravidin-magnetic beads (Fisher Scientific, Cat. No. 11864143. The following day, beads were washed 3 times in ice-cold PBS + 1% Triton X100 and twice with ice-cold PBS only. Samples were then subjected to SDS PAGE and Western blotting probing with antibodies.

### Protein lysate preparation, SDS-PAGE and Western blotting

Cultured astrocytes were washed once with cold PBS and lysed in cold RIPA buffer, supplemented with protease inhibitors (Merck, Calbiochem set III, Cat. No. 539134). Cell lysates were centrifuged at 20,000 g and supernatant was transferred to a tube containing Roti load I SDS sample buffer. On average 15-30 μg of protein was loaded on an SDS-PAGE gel. Western blot analysis was performed as previously described (Schrotter et al., 2016). Quantification of band densities was performed using FIJI. The area of the band and the mean grey value were measured to obtain relative density. For relative quantifications, measurements were normalized to loading control.

### Antibody feeding assay

To visualize internalized active β1-integrin, astrocytes were first starved (DMEM without serum) for one hour and subsequently incubated with the 9EG7 antibody (BD Bioscience, 1:20 in DMEM) on ice for 1 h. After three washes with cold complete medium, cells were incubated for 30 min at 37°C and 5% CO_2_. Astrocytes were washed several times in PBS and then surface antibody was stripped with ice-cold acetic acid pH3 (0.5 M NaCl, 0.5% Acetic Acid in ddH2O) followed by several washes in PBS before fixation.

### *In vivo* stab wound

Mice were anaesthetized with a mixture of ketamine (100 mg/kg) and xylazine (10 mg/kg), and head-fixed on a stereotactic frame (Kopf Stereotax). Throughout the operation, body temperature was maintained at 36-37°C, using a heating blanket. Full anesthesia of the mice was verified throughout the procedure by carefully checking breathing and reflexes using the pinching toe method. For surgery, an incision was created in the scalp and a small craniotomy was drilled above M1 motor cortex (bregma: −1 mm; lateral: 1 mm). An injection needle (Hamilton, gauge 33 was carefully inserted into the motor cortex and moved up and down three times (0.8 mm). The needle was removed and the scalp was sutured. Metamizol (5 mg/ml) was added to drinking water as analgesics until sacrifice. After surgery, mice were replaced in their cage, and kept warm during wake-up and recovery on a heating plate at 37°C. Stab wound outcomes were analyzed 7 or 30 days later by IHC and confocal microscopy.

### Immunohistochemistry and tissue clearance

Seven or 30 days after stab wound injury, mice were sedated with isoflurane, perfused with 4 % formaldehyde and sacrificed. Coronal sections (60 μm diameter) were obtained, permeabilized with 1% Triton-X100 in PBS and blocked in 5% BSA. In a first step, sections were labelled with IHC for GFAP to label reactive astrocytes and identify lesion sites with adjacent glial scars. GFAP-positive sections were stained for additional marker proteins and subsequently fixed in 4% formaldehyde for 1 h at 4°C. Tissue clearance was obtained using ScaleA2 (4 M UREA, 10% (wt/vol) glycerol, 0.1% (wt/vol) Triton X-100 and 0.1x PBS) as described previously (Murk et al., 2013). Transparent slices were mounted on glass slides using Mowiol with 4 M UREA and analyzed using a Nikon A1 confocal microscope with a 40x objective (N.A. 1.3, working distance 240 μm) and automated scanning stage. For visualization of RAB8 tubules *in vivo*, animals were sacrificed as described above and perfused with 3.7% formaldehyde in cytoskeleton-preservation buffer (25 mM HEPES, 60 mM PIPES, 10 mM EGTA, 2 mM MgCl_2_, pH 7.4). Slices were permeabilized with 0.02% Triton X100 overnight and the stained as described above in cytoskeleton-preservation buffer. For visualization of RAB8 tubules *in vivo*, no tissue clearing was performed and cells were mounted on glass slides using Mowiol without UREA.

### Purification of recombinant GST-hSLP4A and isolation of endogenous GTP-bound RAB8 from astrocytes

Recombinant GST-hSLP4A was expressed in and purified from BL21 Rosetta DE3 *E.coli* (Merck, Ca. No. 70954). 5ml of saturated pDEST15 hSlp4-a-transformed BL21 Rosetta DE3 *E.coli* culture were diluted in 500 ml 2x YT (Sigma Aldrich, Cat. No. Y2377-250G) with Ampicillin (Sigma-Aldrich, Cat. No. A9518) under shaking (230 rpm) for 3h at 37°C. Protein expression was induced by 1 mM isopropyl β-d-1-thiogalactopyranoside (IPTG, Sigma-Aldrich, Cat. No. I6758) and occurred overnight at room temperature under shaking. Bacteria were harvested by centrifugation (Ja10 rotor, 5000 rcf for 15 min at 4°C) and stored at −80° C. Purifcation of GST-hSLP4A was performed directly before the isolation of GTP-bound RAB8 from astrocytes: Bacteria were resuspended in (50mM HEPES, 250mM NaCl, 10% glycerol plus 1% Triton X-100, supplemented with protease inhibitors (Merck, Calbiochem set III, Cat. No. 539134) and subsequently sonicated. Lysates were cleared by centrifugation (JA20 rotor, 18000 rpm, 25 min at 4°C). Glutathione sepharose beads (Sigma-Aldrich, Cat. No. GE17-0756-01) were washed three times in HTG buffer (1% Triton, X-100, 25mm HEPES, 150 mM NaCl, 10% glycerol). Bacterial lysates were incubated with glutathione sepharose beads for one hour on a rotating wheel at 4°C. Beads were washed three times with ice-cold lysis buffer (20mM HEPES, 150mM NaCl, 0.5% Triton-X 100). Subsequently, beads were incubated with astrocyte lysate buffer supplemented with protease inhibitors (Merck, Calbiochem set III, Cat. No. 539134) for one hour at 4°C on a rotating wheel. After four times of washing with lysis buffer with protease inhibitors, samples were subjected to SDS-PAGE and Western blotting.

### Electron Microscopy

Brain slices, fixed, permeabilized and labelled for GFAP, were cryoprotected stepwise in 0.1 M sodium phosphate buffer pH 7.4 (PB) supplemented with increasing concentrations of glycerol [10–20–30% (v/v)] and left overnight in 30% glycerol in PBS at 4°C. The tissue was frozen by plunging into hexane (Carl Roth) at −70°C. Samples were transferred into cold methanol (−90°C) in a freeze-substitution chamber (Leica EM AFS). Methanol was replaced three times before the specimens were immersed overnight in anhydrous methanol at −90°C, containing 2% (w/v) uranyl acetate. After rinsing several times with methanol, the temperature was gradually raised to −50°C and left overnight at −50°C. Tissue was then infiltrated with a mix of Lowicryl HM20 resin (Polysciences) and methanol (1:2; 1:1; 2:1, 1 h each) and left in pure resin overnight at −50°C. Samples were transferred to flat embedding molds containing freshly prepared resin at −50°C. UV polymerization was started at −50°C (overnight) and then continued for 4 d at temperatures gradually increasing from −50°C to −20°C (24 h) and finally to +20°C (24 h). Ultrathin sections (70 nm) were mounted on 200-mesh formvar-coated nickel grids (Plano). Images were acquired using a Zeiss EM 900 equipped with a digital camera (Proscan 1K Slow-Scan CCD-Camera).

### Data availability statement

All data generated or analyzed during this study are included in this published article (and its supplementary information files). Additional data are available from the authors upon request.

Cartoons and schemes were created with BioRender.com

