## Supplementary Data for "Drebrin mediates scar formation and astrocyte reactivity during brain injury by inducing RAB8 tubular endosomes"

**Supplementary Information**

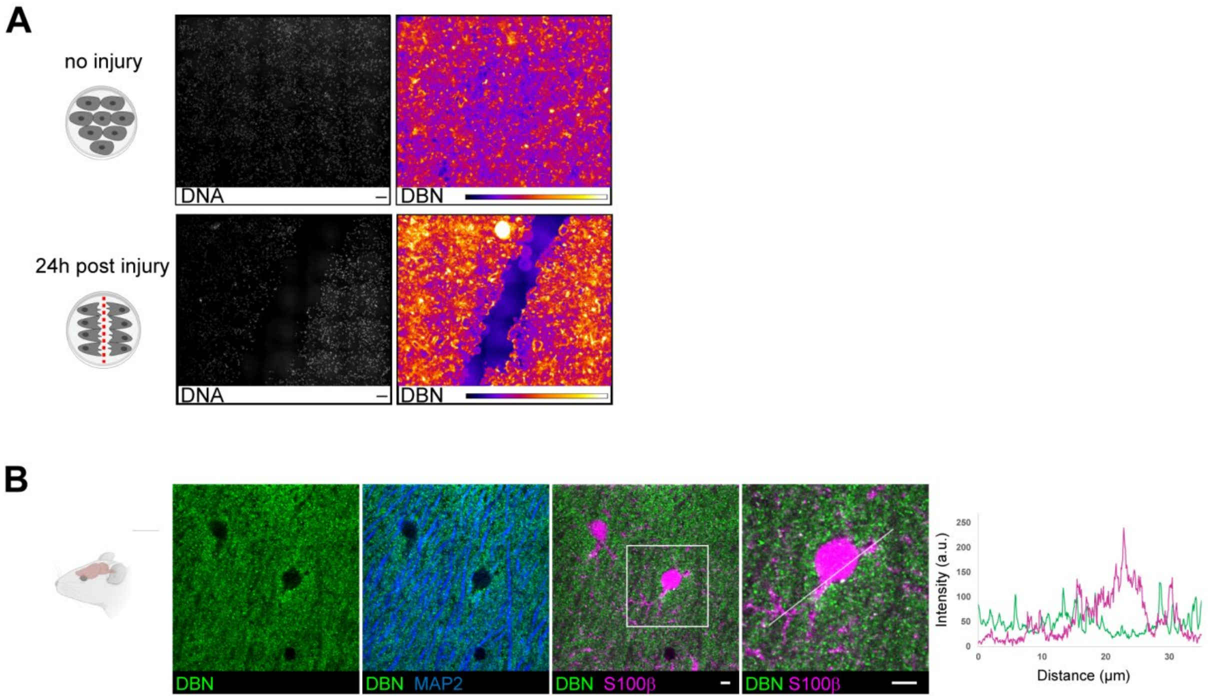


**Supplementary Figure 1: Injury-dependent upregulation of DBN in cultured astrocytes.** (A) Mosaic scan of 25 fields of view from confluent astrocyte cultures labeled with anti-DBN, without or 24 h after mechanical injury. Relative DBN fluorescence is displayed as heatmap. Scale bars: 100 µm. (B) IHC of uninjured or injured mouse brain. Upper panel shows a strong labeling of DBN around MAP2+ dendrites and no DBN labeling in S100ß+ Astrocytes in the uninjured mouse brain. Line-scan through an S100ß+ astrocyte shows little overlap of DBN and S100ß signal.

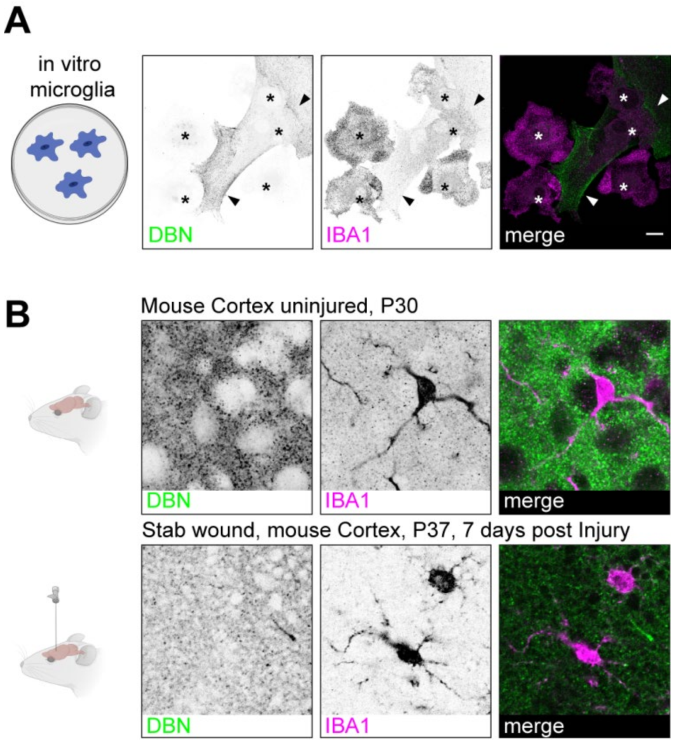


**Supplementary Figure S2: DBN is not expressed in microglia.** (A) Microglia-enriched cultures were labeled for DBN (green) and the microglia marker IBA1 (magenta). IBA1+ microglia are negative for DBN (asterisks), whilst neighboring astrocytes (arrowheads) are DBN+. Scale bars: 10 µm. (B) IHC of DBN (green) and IBA1 (magenta) in the uninjured cortex of P30 WT mice or 7 days post injury. Scale bars: 10 µm.

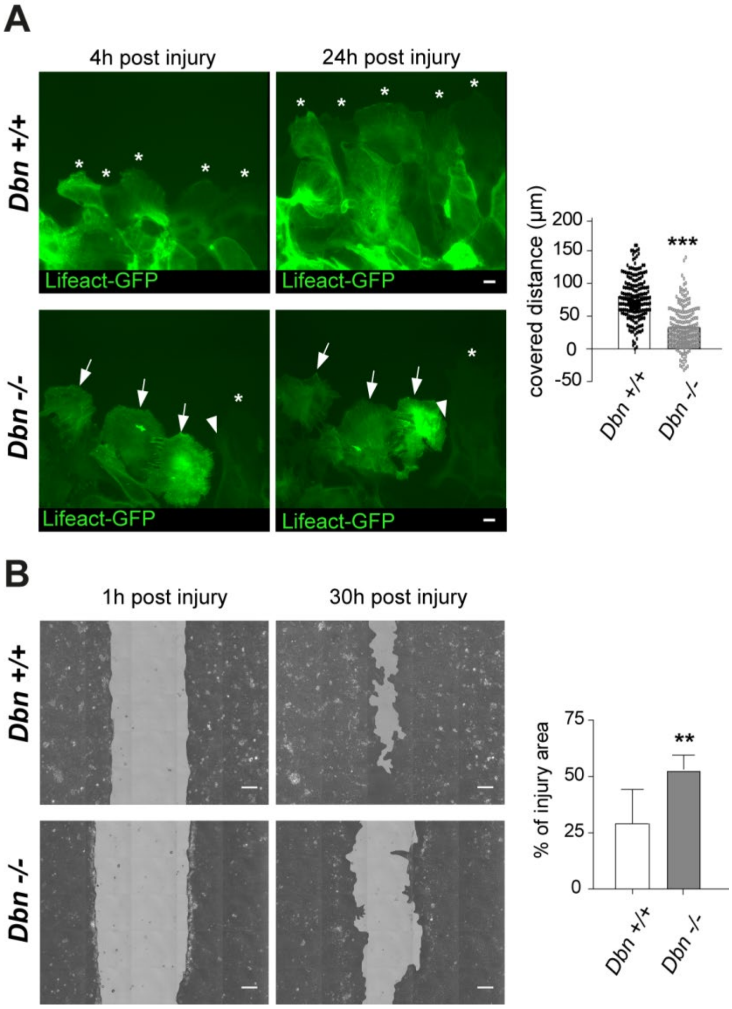


**Supplementary Figure S3DBN-loss perturbs the coordinated outgrowth of cultured astrocytes after scratch injury.** (A) WT and *Dbn*$^{-/-}$ astrocytes expressing Lifeact-GFP, at 4 h and 24 h after mechanical injury *in vitro*. Asterisks indicate cells with extending processes in a persistent manner towards injury. Arrows show moving cells with erratic migratory behavior. The arrowhead show retracting astrocytes. Scale bars: 10 µm. Bar diagram shows quantification of distances covered by WT and *Dbn*$^{-/-}$ astrocytes; n=180 cells per group obtained from three independent experiments, *** $P < 0.001$ (Students unpaired t-test). (B) Live imaging of the overall wound closure of cultured WT and *Dbn*$^{-/-}$ astrocytes after 1 and 30 h. Scale bars: 100 µm. Bar diagram shows quantification of wound size 30 h after injuring the astrocyte monolayers; n=4, ** P<0.01 (Students unpaired t-test).

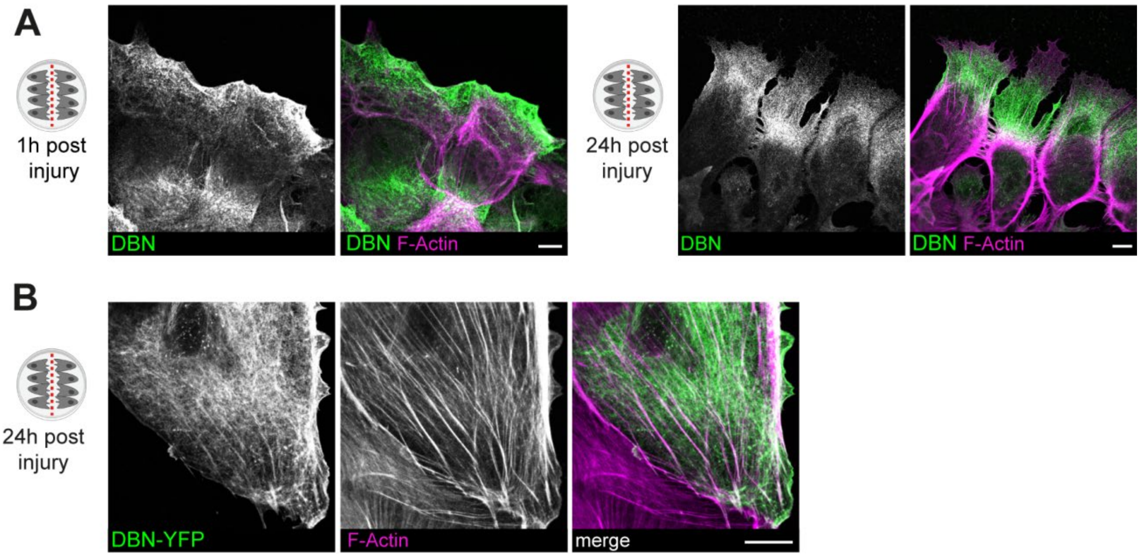


**Supplementary Figure S4: Localization of DBN in cultured astrocytes during scratch injury.** (A) DBN and F-Actin labelling in cultured astrocyte during scratch injury. 1h post injury, DBN localizes to the rear and leading edge of injured astrocytes and shows little co-localization with the most prominent actin filaments. 24h after injury, DBN is detected between the leading edge and the actin-rich cell body, showing again little co-localization with typical actin fibers. Scale bars: 10µm. (B) Localization of DBN-YFP in vesicular and tubular structures and, partially, on actin fibers. Scale bar: 10µm.

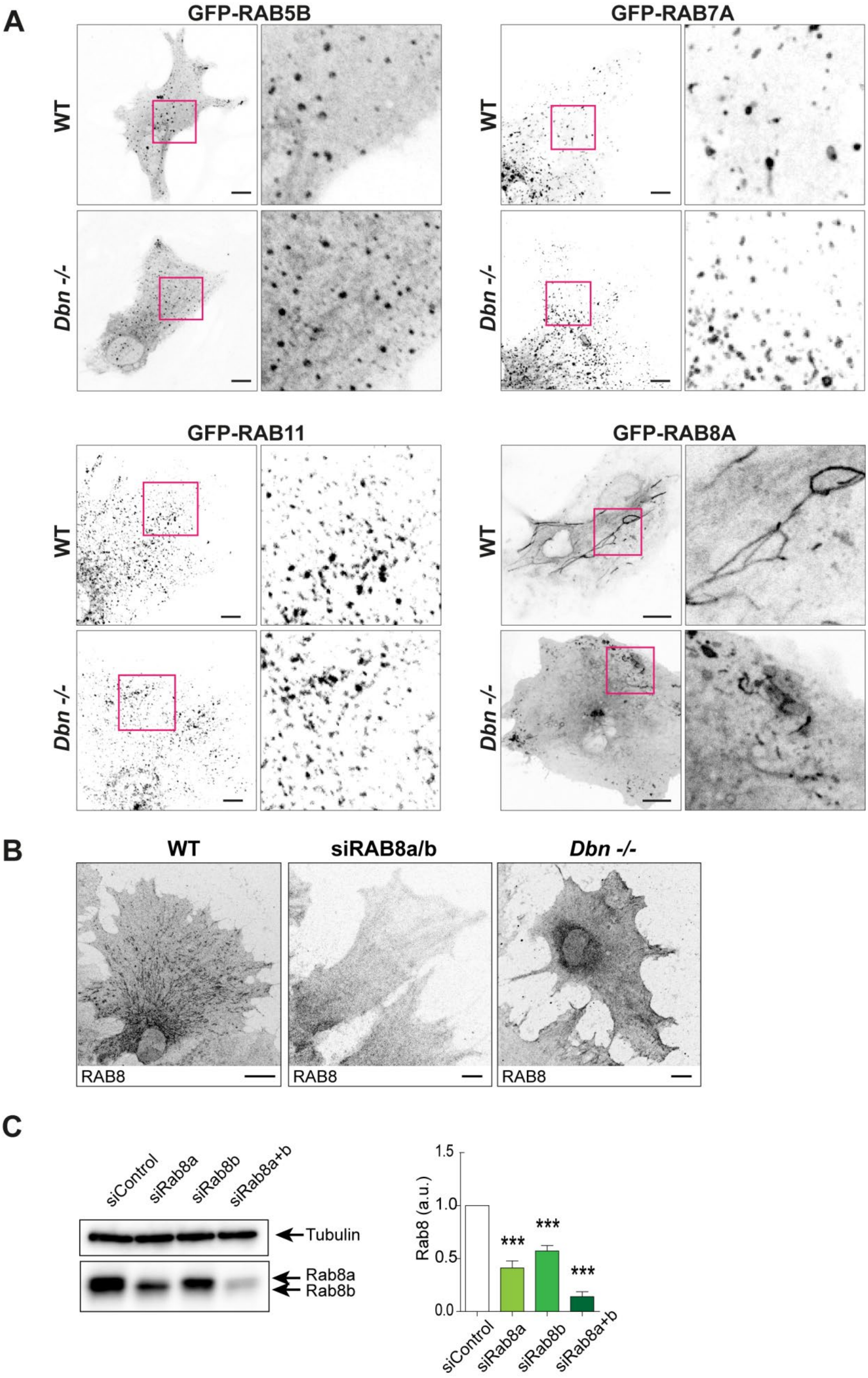


**Supplementary Figure S5: Localization and specificity RAB-GTPases in astrocytes during injury.** (A) Expression of different GFP-tagged RAB GTPases in cultured WT and *Dbn*$^{-/-}$ astrocytes. No major differences between genotypes could be detected for RAB5b, 7A and 11; RAB8A formed tubular structures in WT but not in *Dbn*$^{-/-}$ astrocytes. (B) WT astrocytes, WT astrocytes transfected RAB8A+

siRNA and *Dbn*$^{-/-}$ astrocytes; demonstrating the specificity of the antibody used. (C) Western blot of RAB8 levels using a pan RAB8 antibody in lysates derived from WT astrocytes transfected with control siRNA (siControl), Rab8a specific siRNA (siRab8a) and/or Rab8b specific siRNA (siRab8b). Tubulin immunoreactivity serves as loading control. Bar diagram shows quantification of RAB8 levels from siRNA transfected astrocytes from corresponding western blots) (n=3, ***P<0.001, Students unpaired t-test).

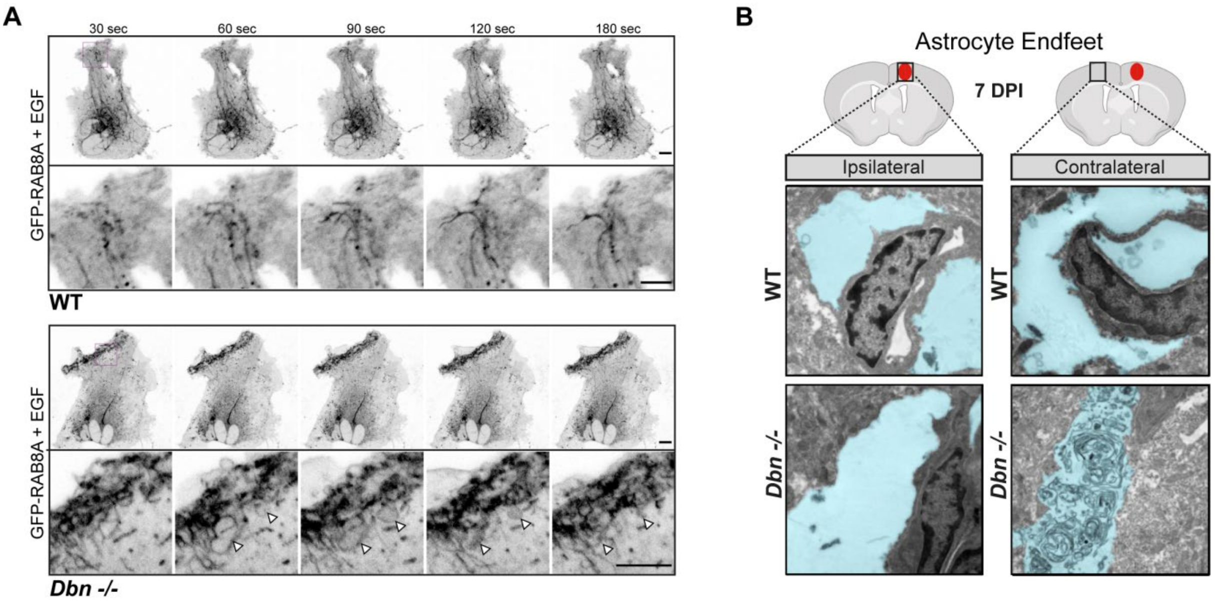


**Supplementary Figure 6: DBN-loss results in accumulation of membranes in astrocytes.** (A) WT and *Dbn*$^{-/-}$ astrocytes expressing GFP-RAB8A after starvation and brief stimulation with EGF (see Movies 4 and 5). Scale bars: 10 µm.(B) TEM images showing accumulation of multilamellar bodies in endfeet of astrocytes in *Dbn*$^{-/-}$ brains, but not in endfeet of astrocytes in WT brains after injury *in vivo* 7 DPI. Astrocyte endfeet are shaded in blue. Scale bars: 500 nm.

**Movie 1**

Live imaging of cultured WT (left) and *Dbn*$^{-/-}$ (right) astrocytes expressing Lifeact-GFP after mechanical injury, corresponding to Supplementary Figure 3. Video acquisition began 4 h and ended 24 h after mechanical injury.

**Movie 2**

Live imaging of cultured WT (left) and *Dbn*$^{-/-}$ (right) astrocytes expressing GFP-RAB8A in conjunction with the parallel visualization of our automated and quantitative tubule detection algorithm. Video recording started 24 hours after injury. Tubules were recorded for 30 min.

**Movie 3**

Live imaging of originally GFP-RAB8A tubule deprived *Dbn*$^{-/-}$ astrocytes mostly after adding Cytochalasin D (left), CK-666 (center) and SMIFH2 (right). Video recording started right after adding the reagents. The effects of the inhibitors were followed for 30 min.

**Movie 4**

Time-lapse video of cultured WT (left) and *Dbn*$^{-/-}$ (right) astrocytes during EGF-stimulation. After 24 h injury, cells were starved for 2h and subsequently treated with EGF. Video recording started immediately after adding EGF. The effect of EGF was followed for 10 min.

**Movie 5**

Magnifications on leading edges of EGF-treated WT (left) and *Dbn*$^{-/-}$ (right) astrocytes according to Movie 4. Note that long extending tubules are formed in *Dbn*$^{+/+}$ astrocytes via smaller particles. EGF treatment leads to membrane accumulation in *Dbn*$^{-/-}$ astrocytes with short tubules and instable vacuole-like structures. Yellow asterisks indicate vacuole-like compartments throughout their lifetime in the *Dbn*$^{-/-}$ time lapse video.

**Movie 6**

Time-lapse video of GFP-RAB8A and microtubules at the leading edge of a *Dbn*$^{-/-}$ astrocyte before and after adding the Arp2/3 inhibitor CK-666, corresponding to Figure 4E. Video recording began 5 min before adding CK-666. The effect of CK-666 was followed for 5 min.

Code for the Analysis of RAB8-tubules:

---

```
/*this codes uses a macro previously published to analyze tubules in cardiomyocytes (Pasqualin et al., 2014). It runs the macro on several images (frames) so that we can analyze the tubules over time and it makes a movie of the skeletonized images. In addition, all results are saved into one .txt file in their order of appearance.*/

//global variables
        var path= " Data_path";
        var name= "picture_name.tif"
//creating 1 table
        table="[Combined Results]";
        run("Table...", "name="+table+" width=450 height=500");
        results_merge=""; //variable that will be filled with all results
//preparing the image, zstack and 8 bit
        run("Z Project...", "start=1 stop=5 projection=[Max Intensity] all");
        run("8-bit");
//code for the skeleton analysis and saving the result files
        Stack.getDimensions(width, height, channels, slices, frames);
        for (i=1; i<frames+1; i++){
          Stack.setFrame(i);
                index=i;
```

```
                        run("Duplicate...", " ");
                        run("Subtract Background...", "rolling=15");
                        run("Enhance Local Contrast (CLAHE)", "blocksize=20 histogram=256 maximum=2
mask=*None*");
                        run("Smooth");
                        run("8-bit");
            //run("Statistical Region Merging", "q=100 showaverages");
                        setThreshold(110, 255);
                        run("Convert to Mask");
                        run("Skeletonize (2D/3D)");
                        saveAs("Tiff", path+"\\frames\\frame_Skeleton"+ index);
                        run("Analyze Skeleton (2D/3D)", "prune=none show original_image=test_xy.tif");
                        run("Summarize");
                        String.copyResults;//these 3 lines are for the table
             resultati=String.paste;
             results_merge=results_merge+resultati;
            selectWindow("Results");
                        saveAs("Results", path+"\\Results\\Results1\\frame_Skeleton_Results"+
index+".xls");
                        selectWindow("Results");
                        selectWindow("Branch information");
                        run("Summarize");
            saveAs("Measurements",path+"\\Results\\branch_information\\_Skeleton_Results_branch"
+ index+".xls");
                        selectWindow("Results");
                        run("Close");
                        selectWindow("_Skeleton_Results_branch"+index+".xls");
                        run("Close");
                        selectWindow("frame_Skeleton"+index+".tif");
                        run("Close");
                        selectWindow("Tagged skeleton");
                        run("Close");
            //close();
                        selectWindow(name);
```

```
            print(i);
        }
            print(table,String.getResultsHeadings);
            print(table,results_merge); //this table contains all results
//code for making a movie of the single skeleton images
            open(path+"\\frames\\frame_Skeleton1.tif")
            dir = getDirectory(path+"\\frames");
            close();
            list = getFileList(dir);
            for (i=0; i<list.length; i++){
              if (File.isDirectory(dir+list[i])){}
              else
             open(dir+list[i]);
            }
            run("Images to Stack", "name=Stack title=[] use");
            saveAs(path+"\\frames\\SkeletonMovie.tif")
close();
```
